# Demonstrating the ability of GABAergic cells in the zona incerta to modulate motivation

**DOI:** 10.64898/2026.05.10.724142

**Authors:** Laura Korobkova, Brian G Dias

**Affiliations:** Neuroscience Graduate Program, University of Southern California, Los Angeles, CA; Developmental Neuroscience and Neurogenetics Program, The Saban Research Institute, Los Angeles, CA; Division of Endocrinology, Diabetes and Metabolism, Children’s Hospital Los Angeles, Los Angeles, CA; Division of Neurology, Children’s Hospital Los Angeles, Los Angeles, CA; Department of Pediatrics, Keck School of Medicine of USC, Los Angeles, CA; Department of Neurology, Keck School of Medicine of USC, Los Angeles, CA

## Abstract

Motivation to engage in goal-directed actions is crucial for survival and well-being. Dopaminergic and serotonergic systems have been the focus of efforts to understand neurobiological etiologies of normative and disrupted motivation. Understanding how the brain incorporates salient sensory cues into motivational drive outside of these neuromodulatory systems is less appreciated. We posited that given their afferent and efferent connections, GABAergic cells in the zona incerta (ZI) are ideally positioned to perform this function. Combining behavioral tasks in mice with chemogenetics we show that GABAergic ZI neurons are capable of modulating effort-based motivation and that chemogenetic activation of this cell population can rescue motivational deficits induced by chronic stress - without affecting attention, memory, locomotion, or appetite. Next, using fiber photometry we report that GABAergic cells in the ZI robustly respond to sensory cues and their response to a cue increases as reward-predictive associations are formed. Inhibiting GABAergic cells in the ZI did not abolish cue-associated motivational behavior while cue-locked optogenetic activation of these cells robustly enhanced cue-associated motivated behavior demonstrating that these cells are not necessary but sufficient to allow for sensory cues to influence action selection. These findings establish GABAergic cells in the ZI as modulators of motivation and afford us a neuroanatomical hub that could be targeted to potentially remedy treatment-resistant deficits in motivation.

## INTRODUCTION

Engaging in goal-directed behavior like seeking out food or social interaction is an important dimension of general well-being. Operationally, motivation to engage in, plan and execute a task is at the core of goal-directed behavior(Salamone & Correa, 2012). Regulated by neural circuits these components of motivation enable organisms to exert effort toward desired outcomes(Schultz, 2015). Disruptions in motivation are central to numerous neuropsychiatric conditions(Nestler & Carlezon, 2006), and the reduced pursuit of rewarding actions is exacerbated by environmental factors like exposure to stress(Russo & Nestler, 2013). Most efforts to understand neurobiological mechanisms that underlie normative motivation and remedy stress-induced disruptions of motivation have focused on mesolimbic and hypothalamic dopaminergic systems in the brain(Baik, 2020; Hollon et al., 2015; Nieh et al., 2016; Stanton et al., 2019). Despite the progress made in these neurobiological arenas, therapeutic strategies targeting these systems frequently fail in many individuals(Al-harbi, 2012; Berlim & Turecki, 2007). Additionally, most studies of motivation pay very little attention to sensory cues that become associated with rewards and then acquire the ability to directly evoke motivational drive and energize goal-directed actions(Berridge & Robinson, 2016; Cartoni et al., 2016; Wolff & Saunders, 2024). Consequently, the circuits that integrate sensory input into motivational drive remain poorly defined. This gap limits our ability to fully map the circuits underlying adaptive motivational control and develop more effective interventions to rescue deficits in motivation. With the goal of not only expanding cellular substrates that could eventually serve as therapeutic targets for stress-induced deficits in motivation but also ensuring that sensory cues are included in such contextualization of motivation, in this study, we sought to determine the contribution of GABAergic cells in the sub-thalamic zona incerta (ZI) to motivation.

The predominantly GABAergic ZI is extensively connected to sensorimotor, limbic, and basal ganglia structures(Arena et al., 2024; Wang et al., 2020). Through this connectivity, GABAergic cells in the ZI integrate incoming sensory information with internal states and motor outputs, and have been implicated in arousal, attention, anxiety, feeding, and reward pursuit(Ahmadlou et al., 2021; Chou et al., 2018; Zhang & van den Pol, 2017; Zhao et al., 2019). These findings make GABAergic cells in the ZI perfect neuroanatomical substrates that could influence normative motivation, remedy stress-induced deficits in motivation and incorporate sensory cues into the drive to be engaged in tasks. Using a combination of chemogenetics, fiber photometry, operant as well as Pavlovian behavioral assays and projection-targeted neuronal manipulations in mice, we identified GABAergic cells in the ZI as important modulators of normative and stress-induced deficits in motivation while dynamically responding to reward-related sensory stimuli and incorporating these stimuli into goal-directed action. Our findings reveal a previously unrecognized role for GABAergic cells in the ZI in integrating sensory and reward information to shape goal-directed action.

## RESULTS

### Chemogenetic manipulation of GABAergic neurons in the ZI bidirectionally modulates motivation

To test whether GABAergic neurons in the ZI contribute to motivational drive, we expressed inhibitory (hM4DGi) or excitatory (hM3DGq) DREADDs in ZI GABAergic neurons by bilaterally injecting Cre-dependent AAVs (AAV-hSyn-DIO-DREADD-mCherry or AAV-hSyn-DIO-eGFP) into the ZI of vGAT-CRE mice **(Fig. 1A and 1B)**. Following surgical recovery and operant training on fixed-ratio schedules, motivation in the mice was tested on a progressive ratio (PR) schedule **(Fig. 1C)**. In this well-established assay of motivational vigor, the response requirement increases exponentially with each successive reinforcer, and the breakpoint (the highest ratio completed) serves as the primary index of motivation.

**Figure 1.**
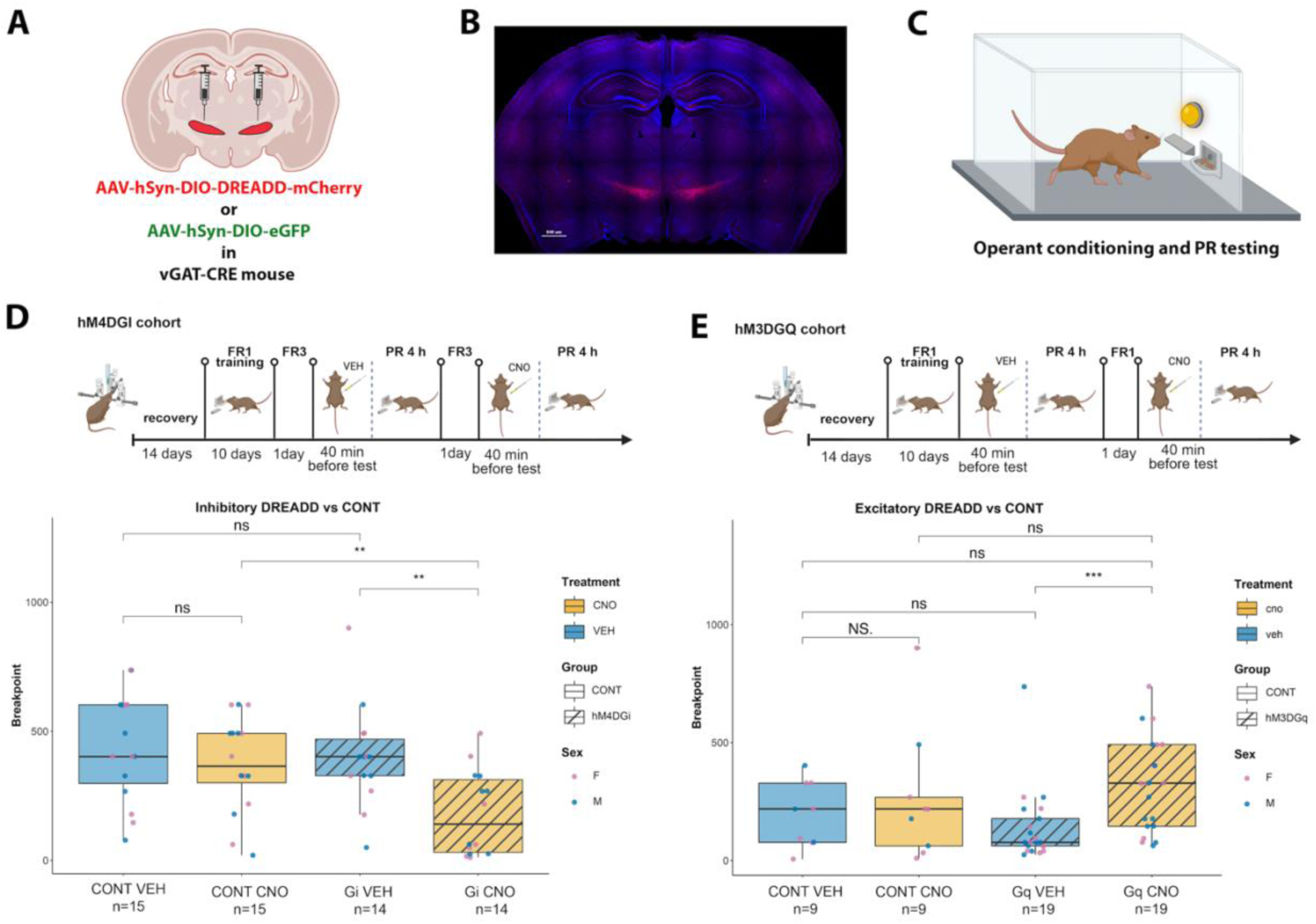
Chemogenetic manipulation of GABAergic neurons in the ZI bidirectionally modulates motivation. **(A)** Schematic of experiment: CRE-dependent inhibitory (hM4DGi) or excitatory (hM3DGq) DREADDs or GFP control were bilaterally injected into the ZI of vGAT-CRE mice. **(B)** Representative histological image showing DREADD expression in the ZI. **(C)** Schematic of the operant conditioning box. **(D)** Experimental timeline and Progressive Ratio test results for the inhibitory DREADD (hM4DGi) cohort. Chemogenetic inhibition of GABAergic neurons in the ZI significantly reduced breakpoints following CNO administration relative to vehicle and relative to GFP controls. GFP controls showed no treatment-dependent change. **(E)** Experimental timeline and Progressive Ratio test results for the excitatory DREADD (hM3DGq) cohort. Chemogenetic activation of GABAergic neurons in the ZI significantly increased breakpoints following CNO administration compared to vehicle, with no effect in GFP controls. Whisker plots show median and interquartile range; individual data points are colored by sex. **p < 0.01, *p < 0.05, ns = not significant. Note that the hM4DGi (D) and hM3DGq (E) cohorts were independent experiments trained on FR3 and FR1 schedules, respectively; absolute breakpoints are therefore not comparable across panels and no cross-experiment statistical comparison is made.

The inhibitory and excitatory cohorts were run as two independent experiments and were trained on different fixed-ratio schedules prior to progressive ratio testing: mice in the hM4DGi cohort were trained on an FR3 schedule, whereas mice in the hM3DGq cohort were trained on an FR1 schedule. Because prior fixed-ratio training history shifts the absolute level of responding on a subsequent progressive ratio schedule, baseline breakpoints are not directly comparable across the two cohorts, and the GFP controls of the hM4DGi experiment accordingly show higher absolute breakpoints than those of the hM3DGq experiment. We therefore make no statistical comparison between the two experiments; all inferences are drawn within experiment, from the within-subject vehicle-versus-CNO contrast and from the comparison against the concurrently run GFP controls of the same cohort.

In our inhibitory experiment, to account for the repeated-measures design (each animal receiving both vehicle and CNO), a linear mixed-effects model was fitted with treatment, group, and sex as fixed effects and animal as a random intercept. The model revealed a significant main effect of treatment (F(1, 35.9) = 12.17, p = 0.001) and, critically, a significant treatment × group interaction (F(1, 35.9) = 9.60, p = 0.004). No main effect of group (F(1,36.4) = 0.13, p = 0.72) or sex (F(1, 36.4) = 0.63, p = 0.43) was observed, and no interactions involving sex reached significance (all p > 0.28). More specifically, chemogenetic inhibition (hM4DGi group: n=14: 7 males, 7 females) of GABAergic ZI neurons in the ZI after CNO administration significantly reduced breakpoints relative to VEH (vehicle) treatment (W = 158.5, p-value = 0.005498, Cohen’s d = 1.347753) and relative to GFP controls (n=15: 8 males, 7 females) treated with CNO (W = 175, p-value = 0.008851, Cohen’s d = 1.08543) **(Fig. 1D)**. Importantly, this reduction in breakpoint was specific to the Gi DREADD group following CNO administration; GFP controls showed no change in breakpoint between vehicle and CNO sessions (W = 143, p-value = 0.3685), confirming that the effect was not attributable to CNO alone.

In the excitatory DREADD (hM3DGq) experiment, a linear mixed-effects model with treatment, group, and sex as fixed effects and animal as a random intercept revealed a significant main effect of treatment on progressive ratio breakpoint (F(1, 52) = 4.583, p = 0.03699). The treatment × group interaction did not reach significance (F(1, 52) = 1.3613, p = 0.24864), nor did any other main effects or interactions (all p > 0.2). Planned comparisons (Wilcoxon signed-rank test) showed that CNO significantly increased breakpoint relative to vehicle in the hM3DGq group (n=19, 10 males, 9 females) (W = 82.5, p-value = 0.0026, Cohen’s d = 1.026742) whereas GFP controls (n=9, male = 3, female = 6) again showed no treatment-dependent change (W = 40, p-value = 1) **(Fig. 1E)**. Together, these results demonstrate that activity of GABAergic cells in the ZI bidirectionally and causally modulates motivational effort, though the two directions are not equally strong. Inhibition produced a significant treatment × group interaction and a CNO-induced reduction in breakpoint that was significant both against the animals’ own vehicle sessions and against CNO-treated GFP controls. Activation produced a significant main effect of treatment and a significant within-subject vehicle-versus-CNO increase, but CNO-treated hM3DGq mice did not differ significantly from CNO-treated GFP controls. The enhancement of motivation by activating GABAergic ZI neurons in well-trained animals is therefore a weaker and less specific effect than the suppression produced by inhibiting them.

Given that we saw a dramatic reduction in motivation after chemogenetically inhibiting GABAergic neurons in the ZI, we wanted to rule out confounding effects of locomotor capacity, memory, or homeostatic hunger drive. Chemogenetic inhibition of GABAergic neurons in the ZI did not alter total distance traveled in the open field (F(3,37)= 1.082, p-value= 0.369) **(Fig. S1)**, had no effect on preference index in a memory task(F(1,30)= 0.369, p-value=0.57025) **(Fig. S2)**, and did not change food consumption (W = 24.5, p-value = 1) **(Fig. S3)**. These data indicate that the observed changes in breakpoint reflect a specific modulation of motivational state rather than generalized alterations in motor output, cognitive function, or appetitive drive.

### Chemogenetic activation of GABAergic neurons of the ZI rescues chronic stress-induced deficits in motivation

To determine whether activation of GABAergic ZI neurons is sufficient to rescue stress-induced deficits in motivation, we expressed excitatory DREADDs (AAV-hSyn-DIO-hM3DGq-mCherry) or control virus (AAV-hSyn-DIO-eGFP) in GABAergic neurons in the ZI of vGAT-CRE mice and assigned animals to one of three groups: no stress/GFP control (n=7, 3 males, 4 females), stress/GFP (n=6, 2 males, 4 females), and stress/hM3DGq (n=8, 4 males, 4 females) **(Fig. 2A-C)**. Following recovery from surgery, mice in the stress/GFP and stress/hM3DGq groups were exposed to a chronic stress protocol concurrent with operant training on FR schedules **(Fig. 2D)**. Critically, all three groups acquired the operant response at comparable rates, with no significant differences in reinforcements earned across training days, indicating that stress exposure did not impair task learning (F(26) = 1.27, p > 0.05) **(Fig. S4)**.

**Figure 2.**
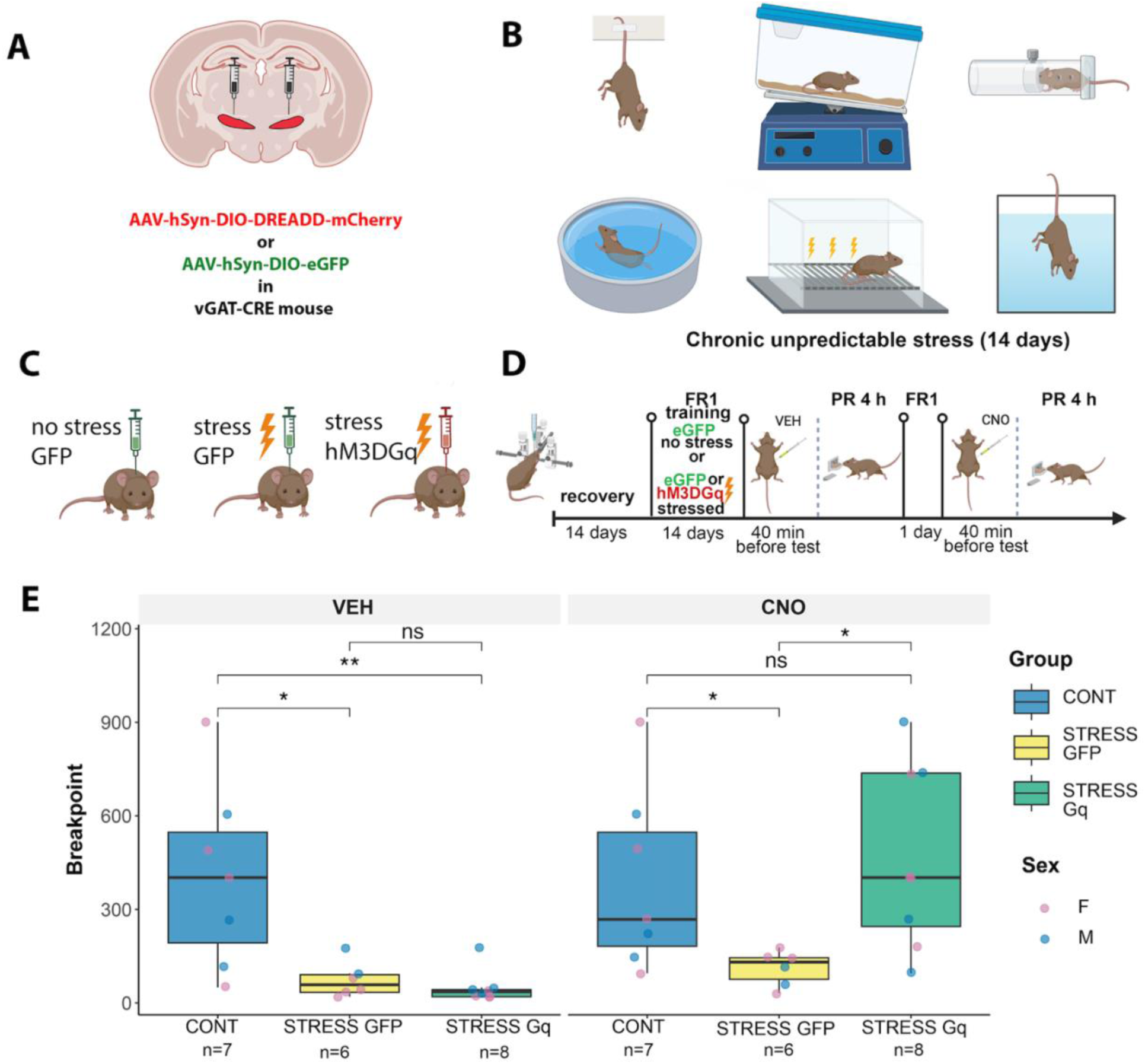
Chemogenetic activation of GABAergic ZI neurons rescues stress-induced deficits in motivation. **(A)** Schematic of viral strategy: excitatory DREADDs (hM3DGq) or GFP control were bilaterally injected into the ZI of vGAT-Cre mice. **(B)** Schematic of the chronic unpredictable stress protocol. (C) Experimental groups: no stress/GFP control, stress/GFP, and stress/hM3DGq. **(D)** Experimental timeline showing concurrent stress exposure and operant training on FR schedules, followed by PR testing under vehicle and CNO conditions. **(E)** Breakpoint data under vehicle (left) and CNO (right) conditions. Under vehicle, both groups exposed to stress showed significantly reduced breakpoints compared to non-stressed controls. Following CNO administration, hM3DGq mice exposed to stress showed breakpoints no longer significantly different from controls. Whisker plots show median and interquartile range; individual data points are colored by sex. **p < 0.01, *p < 0.05, ns = not significant.

A linear mixed-effects model was fitted with treatment (vehicle vs. CNO), group (control, stress/GFP, stress/hM3DGq), and sex as fixed effects and animal as a random intercept. The model revealed a significant main effect of treatment (F(1, 15) = 10.89, p = 0.005) and, critically, a highly significant treatment × group interaction (F(2, 15) =11.93, p < 0.001), indicating that the effect of CNO on breakpoint differed across groups. A trending main effect of group was also observed (F(2, 15) = 2.82, p = 0.091), while sex and all sex-related interactions were non-significant (all p > 0.69). During the PR test under vehicle conditions, mice exposed to stress exhibited significantly lower breakpoints compared to non-stressed controls (stress/GFP vs no stress/GFP: W=38, p-value= 0.01399, Cohen’s d = 1.497416; stress/ hM3DGq vs no stress/GFP: W=53.5, p-value= 0.003628, Cohen’s d = 1.740187), confirming that chronic stress reduced motivational effort **(Fig. 2E, left)**. However, upon CNO administration, hM3DGq mice exposed to stress showed a marked increase in breakpoint that was no longer significantly different from non-stressed controls (W=23.5, p-value= 0.6419), whereas stressed GFP mice still exhibited significant reduction in motivation (W=36, p-value= 0.03726, Cohen’s d = 1.264385) **(Fig. 2E, right)**. These findings demonstrate that chemogenetic activation of GABAergic neurons of the ZI is sufficient to rescue motivation following chronic stress.

### GABAergic neurons of the ZI fire in response to sensory stimuli across multiple modalities

Motivation is not a unitary process but rather a temporally extended sequence of responses: detection of reward-predictive cues, initiation of goal-directed action, execution of instrumental responses, and evaluation of reward outcomes. While our findings established GABAergic neurons in the ZI as regulators of motivated behavior, the specific components of motivation that these neurons govern were left unresolved. The ZI receives convergent sensory input from diverse brain regions. However, whether GABAergic neurons in the ZI respond to moments of intention, action, or sensory events during motivated behavior has not been directly examined. To dissect the contribution of the ZI to distinct phases of motivational processing, we expressed GCaMP in GABAergic neurons in the ZI of vGAT-CRE mice and performed fiber photometry recordings during operant behavior. Real-time calcium dynamics were recorded while animals performed an operant task on a modified FR1 schedule designed to isolate sensory, motor, and reward-related components of motivated behavior by disabling lever retraction after press and delivering reward after a delay **(Fig. 3A)**. This design temporally separates neural responses associated with the instrumental action from those evoked by reward delivery.

**Figure 3.**
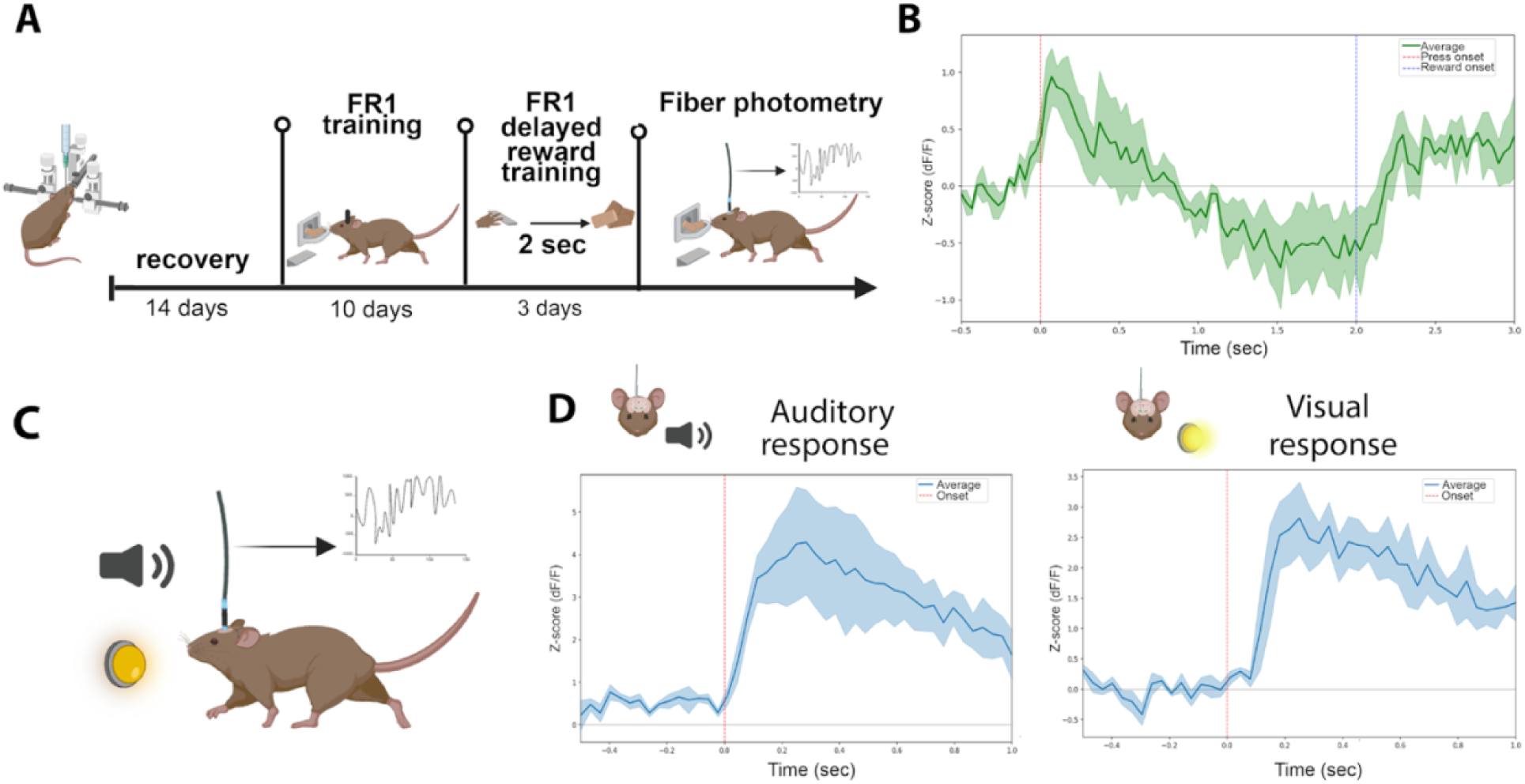
GABAergic neurons in the ZI respond to sensory stimuli across multiple modalities. **(A)** Experimental timeline: vGAT-CRE mice expressing GCaMP in GABAergic neurons in the ZI underwent FR1 operant training followed by a modified FR1 task with delayed reward delivery, during which fiber photometry recordings were performed. **(B)** Mean z-scored calcium signal (dF/F) from GABAergic neurons in the ZI time-locked to lever press and reward delivery during the delayed-reward FR1 task. Per-trial SD baseline z-scoring applied. Shaded regions represent SEM. **(C)** Schematic of multimodal sensory stimulation paradigm in naive mice. (D) Mean z-scored calcium transients in ZI-located GABAergic neurons in response to auditory (left) and visual (right) stimuli, showing significant time-locked responses to both sensory modalities.

During FR1 training, we observed robust calcium transients in GABAergic ZI neurons time-locked to both lever press and reward delivery, with a characteristic two-component response: an initial signal increase upon lever press followed by a sustained activation upon reward retrieval **(Fig. 3B)**.

To assess whether the response of GABAergic ZI neurons was specific to reward delivery, we tested how these neurons respond to neutral sensory stimuli of different modalities. Mice naïve to the operant chamber were exposed to brief auditory and visual stimuli while calcium dynamics were recorded **(Fig. 3C)**. GABAergic ZI neurons exhibited time-locked increases in z-scored ΔF/F in response to these sensory modalities, with responses emerging rapidly after stimulus onset and returning to baseline within approximately one second **(Fig. 3D)**. To determine whether GABAergic ZI neurons respond differently to sensory cues, we compared the waveform of peri-event calcium transients evoked by lever press, an auditory cue and a visual cue **(Fig. S5)** All three event types were analysed on an identical peri-event window (−0.5 to +1.0 s relative to event onset) so that latency and kinetic measures were directly comparable, and decay was quantified over a fixed 0.4 s interval following the peak, Responses differed in magnitude across event types (Kruskal–Wallis, H = 14.59, p = 0.0007): auditory-evoked transients were approximately twice as large as those evoked by lever press (median 1.36 vs 0.74 z, Mann–Whitney U = 0, p = 0.0002) or by the visual cue (1.36 vs 0.68 z, U = 52, p = 0.0016), whereas press- and light-evoked transients were of comparable amplitude (U = 29, p = 0.29). Latency to peak also differed across event types (H = 6.75, p = 0.034), but the difference was carried entirely by the visual cue. Decay was steepest for auditory-evoked transients (H = 7.71, p = 0.021; sound −1.62 z/s vs press −0.53 z/s, U = 54, p = 0.016; vs light −1.05 z/s, U = 8, p = 0.026). When decay was expressed as the fraction of the peak remaining after 0.4 s, an amplitude-independent measure, no difference across event types was detected (H = 0.45,p = 0.80), indicating that the steeper absolute decay of auditory responses reflects their larger amplitude rather than faster kinetics. Individual animal traces (n=3) confirmed the consistency of these responses across subjects **(Fig. S6)**. These data reveal that GABAergic ZI neurons are broadly tuned to sensory input across modalities.

### GABAergic neurons of the Zona Incerta encode the learned motivational valence of sensory cues

Given that GABAergic ZI neurons respond to sensory stimuli and are involved in regulation of motivated behavior, we next asked whether responses to sensory cues are modulated by the learned motivational significance of the cue. To test this, we employed a Pavlovian discrimination paradigm in which one visual cue (CS+; e.g., light on the right) was paired with reward delivery and a second visual cue (CS-; e.g., light on the left) was presented without reward, while recording calcium transients in GABAergic ZI neurons via fiber photometry **(Fig. 4A, 4B)**.

**Figure 4.**
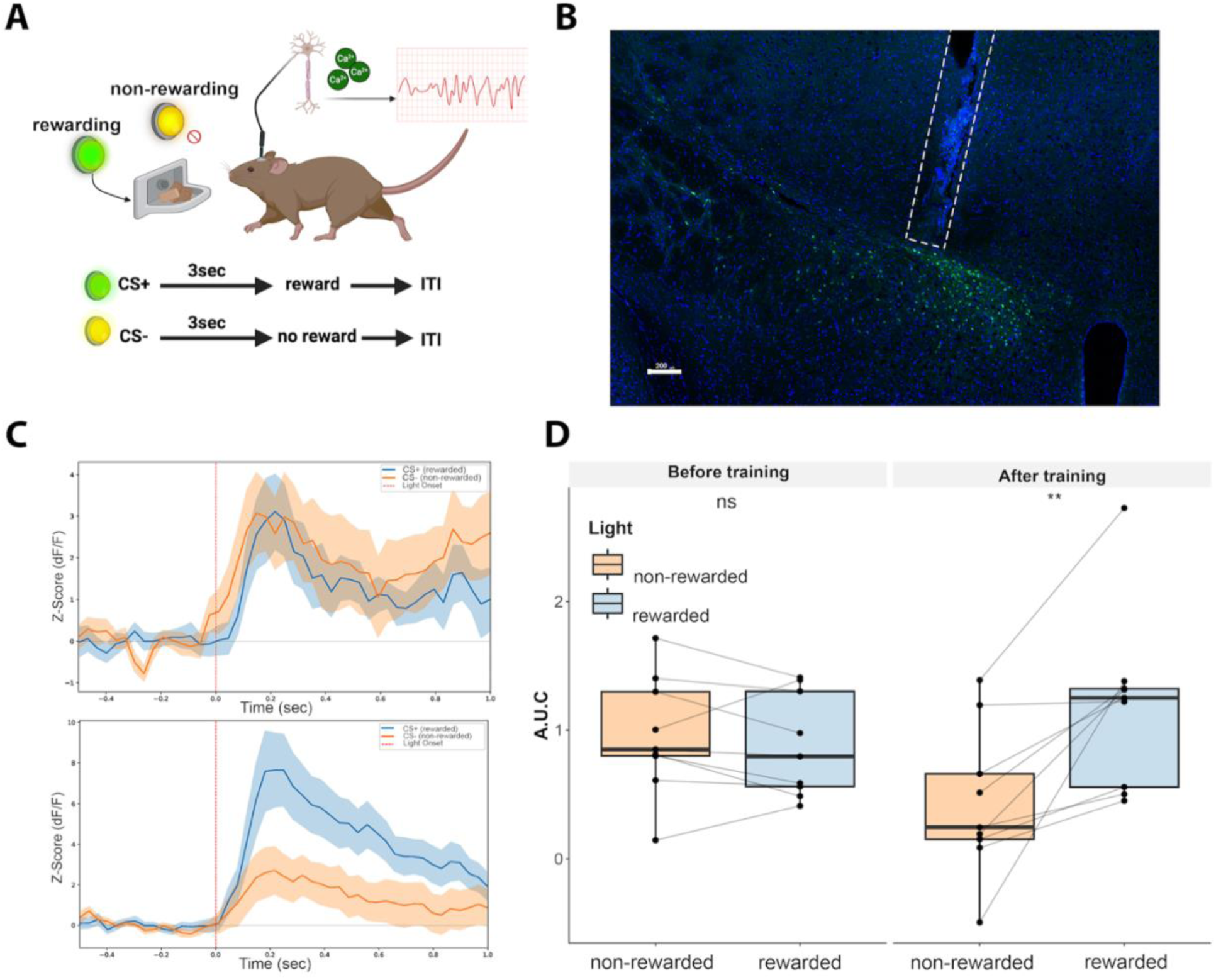
GABAergic ZI neurons encode the learned motivational valence of sensory cues. **(A)** Schematic of Pavlovian discrimination paradigm: one visual cue (CS+) was paired with reward delivery and a second visual cue (CS-) was presented without reward, while calcium transients were recorded via fiber photometry. **(B)** Representative histological image showing DIO-GCaMP expression and fiber placement in the ZI. **(C)** Mean z-scored calcium traces in response to CS+ and CS-before training (top) and after training (bottom). Data represented as mean ± SEM (shaded area). Dashed red line indicates light onset. **(D)** Quantification of calcium responses (area under the curve of z-scored dF/F) to rewarded and non-rewarded cues before and after training (n=9, 5 males, 4 females). After training, responses to the CS+ were significantly greater than to the CS-. Data represented as boxplots (median, interquartile range, whiskers) with individual data points connected across conditions by lines. **p < 0.01, ns = not significant.

GABAergic ZI responses were quantified as the area under the curve (AUC) of z-scored ΔF/F during the 1-second window following CS presentation. A three-way ANOVA examining the effects of CS, time (pre-vs post-training), and sex on GABAergic ZI activity revealed a significant CS × training interaction (F(1, 28) = 5.66, p-value = 0.025), with no significant interactions involving sex (F(1, 28) = 0.482, p-value=0.4932). Before conditioning, GABAergic ZI neurons responded comparably to both cues, consistent with the observed general sensory responsiveness (W=33, p-value = 0.5457) **(Fig. 4C, top)**. However, after Pavlovian training, a striking divergence emerged: the CS+ elicited a larger calcium transient than the CS-(W=65, p-value = 0.03147, Cohen’s d=1.178327) **(Fig. 4C, bottom)**. Quantification of the responses confirmed that, before training, there was no significant difference in GABAergic ZI responses between the rewarded and non-rewarded visual cues, whereas after training, responses of GABAergic neurons of the ZI to the CS+ were significantly greater than those to the CS-**(Fig. 4D)**. Behavioral evidence of successful discrimination was further supported by a progressive decrease in inter-response time (IRT) between light onset and detector entry selectively for the CS+ across training days (LME, p-value < 0.01) **(Fig. S7)**. These results demonstrate that GABAergic ZI neurons do not merely relay sensory information but dynamically update their responses to reflect the acquired motivational valence of environmental cues.

### Inhibition of GABAergic ZI neurons does not suppress reward-associated conditioned responding

Our photometry data revealed that ZI GABAergic neurons encode the learned valence of reward-predictive cues. We therefore asked whether silencing these neurons would disrupt the ability of a conditioned stimulus to evoke goal-directed responding. vGAT-CRE mice expressing inhibitory DIO-DREADDs (Gi) or DIO-GFP control virus in the GABAergic neurons of the ZI underwent Pavlovian conditioning in which an auditory cue (CS+) was paired with reward and a visual cue (CS-) was unreinforced **(Fig. 5A-B)**. Following conditioning during which all groups acquired operant training **(Fig. S8)**, mice were trained on an FR1 operant schedule and subsequently underwent an operant probe test in which the CS+ and CS-were presented in alternating blocks without reinforcement upon CNO condition **(Fig. 5C)**.

**Figure 5.**
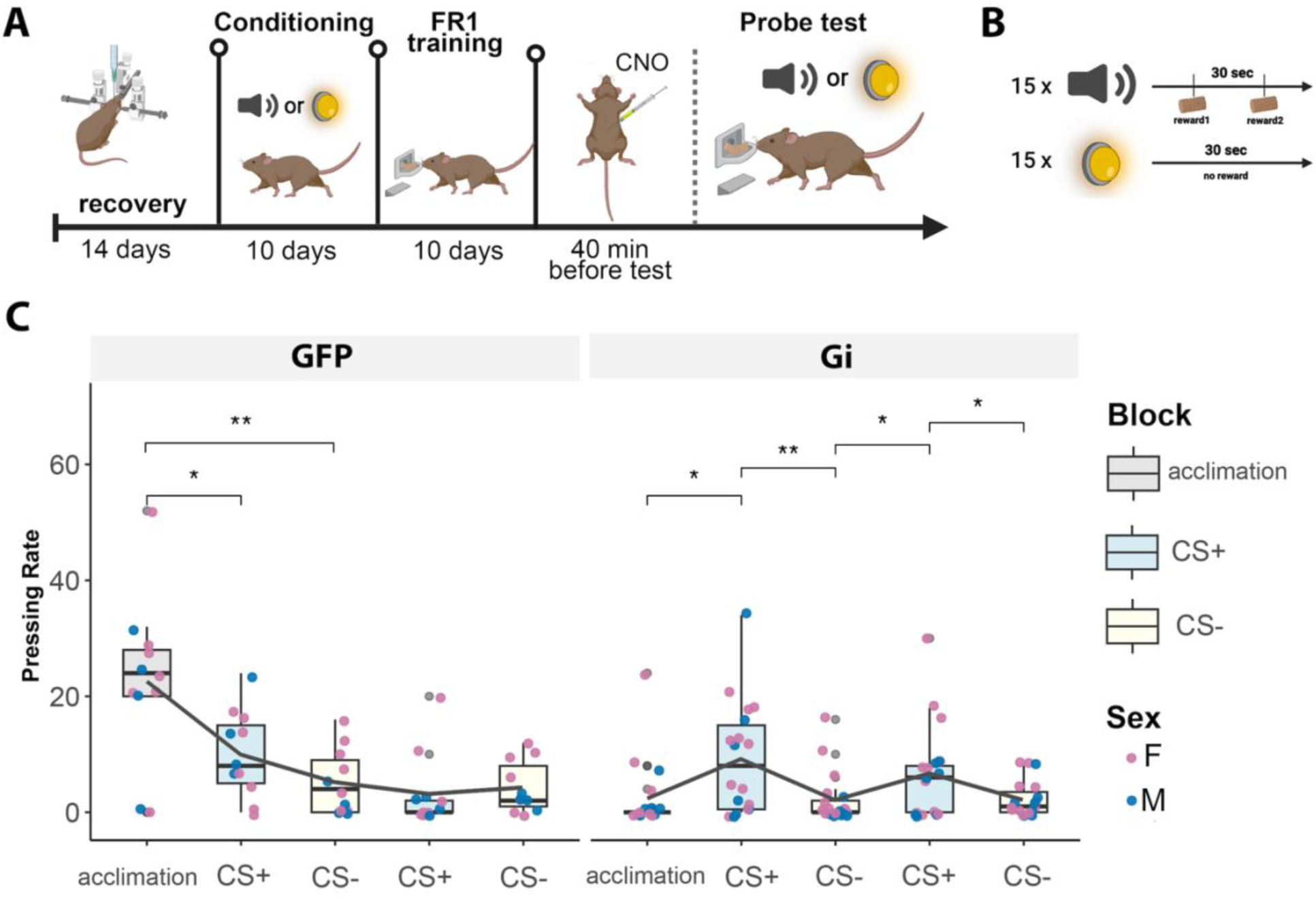
Inhibition of GABAergic ZI neurons does not suppress reward-associated conditioned responding. **(A)** Experimental timeline: vGAT-CRE mice expressing inhibitory DIO-DREADDs (Gi) or DIO-GFP control underwent Pavlovian conditioning, FR1 operant training, and an operant probe test with CS presentations under CNO. **(B)** Pavlovian conditioning design: an auditory cue (CS+) was paired with reward and a visual cue (CS-) was unreinforced. **(C)** Pressing rate during the probe test across alternating CS+, CS-, and acclimation time blocks for GFP controls (left) and Gi DREADD mice (right). GFP controls showed a progressive decline in pressing with no differential response to CS+ versus CS-. Gi DREADD mice showed selective increases in pressing rate during CS+ blocks relative to both acclimation and CS-blocks. (Gi: n=18, 7 males, 11 females; GFP: n=11, 4 males, 7 females). Comparisons tested: acclimation: CS+ (first presentation); acclimation: CS-(first presentation); CS+ (first presentation): CS-(first presentation); CS+ (first presentation): CS-(second presentation); CS+ (second presentation): CS-(second presentation). **p < 0.01, *p < 0.05, Only significant comparisons are shown. Data represented as boxplots (median, interquartile range, whiskers) with individual data points colored by sex and group mean connected by lines.

A linear mixed-effects model examining pressing rate during the probe test under CNO condition, with block, group (Gi vs GFP), and sex as fixed factors and animal as a random effect revealed significant block × group interaction (F(4, 100) = 9.39, p < 0.001), indicating that the two groups responded differently across block types. There were no significant interactions involving sex (all p > 0.46). Post-hoc comparisons (paired Wilcoxon signed-rank tests, averaged across presentations) showed that GFP controls decreased pressing relative to acclimation during both CS+ (V = 4, p = 0.011) and CS− blocks (V = 3, p = 0.009), with no difference between cues (V = 33.5, p = 0.57) **(Fig. 5C)**. In contrast, Gi mice selectively increased pressing during CS+ blocks (CS+ vs acclimation: V = 106.5, p = 0.009; CS− vs acclimation: V = 48, p = 0.50) and pressed more during CS+ than CS− blocks (V = 112.5, p = 0.003). When the acclimation period was excluded from the model, the block × group interaction was no longer significant (F(3,75)=1.24, p=0.3; nor was the cue x group interaction when the wro presentations of each cue were modelled as repeated measures within animal (F(1,27) = 2.39, p = 0.13). Importantly, however, acclimation pressing did not moderate CS+/CS− discrimination (cue × acclimation, F(1, 26) = 0.01, p = 0.93), and within the Gi group the CS+ > CS− difference remained significant when acclimation rate was included as a covariate (F(1, 16) = 10.1, p = 0.006). A model of cue-block pressing including block position and cue identity as fixed effects confirmed that GFP responding declined progressively across the session (β = −1.9 presses per block, p = 0.040), with no effect of cue identity (p = 0.96), indicating a cue-independent decline **(Fig. 5C).** In Gi mice, we observed the opposite effect: no within-session decline (p = 0.43), but a significant increase in pressing during CS+ blocks (β = 5.2 presses, p = 0.004). Thus, discrimination in Gi mice was driven by cue identity, and was attributable neither to their low baseline nor to time in session.

These data indicate that, although chemogenetic inhibition of GABAergic ZI neurons reduces the general motivational drive to work for reward **(Fig. 1D)**, it does not abolish previously established cue–reward associations or the capacity of conditioned stimuli to selectively enhance instrumental responding.

### Temporally precise optogenetic activation of GABAergic neurons of the ZI during reward-predictive cues is sufficient to enhance motivation

To establish a causal link between sensory-evoked ZI activity and motivational drive with temporal precision, we employed optogenetic stimulation of GABAergic ZI neurons time-locked to reward-predictive sensory cues during progressive ratio testing. Following ChR2 (AAV5-Ef1a-DIO hChR2(E123T/T159C)-EYFP) injection and unilateral surgical implantation of an optical fiber above the ZI **(Fig. 6A)**, mice were first trained on FR1 - one lever press gives one reward, then on a contingent FR1 schedule in which a visual cue (light) signaled reward availability **(Fig. 6B)**. During this contingent FR1 schedule, the lever was available to be pressed throughout the entire session, but presses were reinforced with reward only during light-on periods (each 10 seconds long). After acquiring the contingent association, evidenced by progressively higher pressing rates during light-on versus light-off periods **(Fig. 6C)**, mice were tested on a PR schedule for an hour under three counterbalanced conditions: optogenetic stimulation coincident with the light cue, temporally random stimulation, and no stimulation.

**Figure 6.**
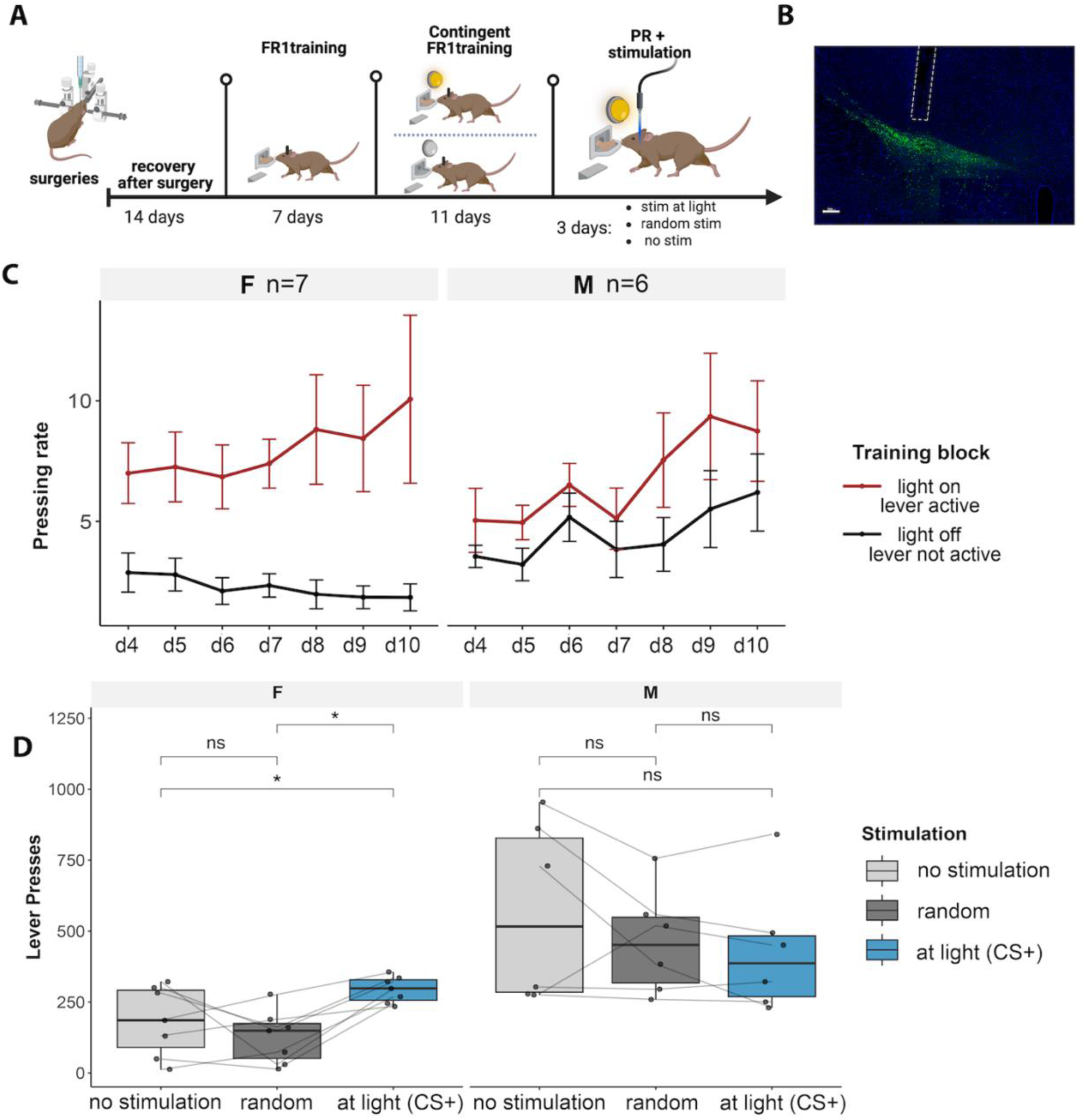
Temporally precise optogenetic activation of GABAergic ZI neurons during reward-predictive cues enhances motivation. **(A)** Representative histological image showing DIO-ChR2 expression and fiber placement in the ZI. **(B)** Experimental timeline: vGAT-Cre mice expressing ChR2 in the ZI underwent FR1 training, contingent FR1 training with a visual cue signaling reward availability, and PR testing under three counterbalanced stimulation conditions. **(C)** Pressing rates during light on (lever active) and light off (lever not active) periods across contingent FR1 training days for females (left) and males (right), showing acquisition of cue-contingent responding**. (D)** Total lever presses on the PR schedule under no stimulation, random stimulation, and stimulation time-locked to the light cue (CS+) for females (left) and males (right). In females, contingent optogenetic stimulation significantly increased lever pressing compared to both random and no stimulation conditions. This effect was not observed in males. Data represented as boxplots (median, interquartile range, whiskers) with individual data points, *p < 0.05, ns = not significant.

In female mice, contingent optogenetic stimulation time-locked to the reward-predictive light cue significantly increased total lever presses on the PR schedule compared to both the no-stimulation and random stimulation conditions (stim at light vs random stim: V = 0, p-value = 0.01563; stim at light vs no stim: V = 2, p-value = 0.04688) **(Fig. 6D, left)**. Notably, random stimulation did not significantly increase lever pressing relative to the no stimulation condition (V = 9, p-value = 0.4688), demonstrating that the enhancement of motivational output required temporal coincidence between GABAergic ZI activation and the sensory cue rather than generalized excitation. To rule out the possibility that these effects were driven by order or practice effects rather than stimulation condition, we reanalyzed the data by testing day rather than stimulation pattern **(Fig. S9)**.

This effect was not observed in male mice, for whom no stimulation condition significantly altered PR performance (V = 10, p-value = 1) **(Fig. 6D, right)**. To explore the basis of this sex difference, we examined behavioral strategies during training. Female mice progressively suppressed responding during light-off periods across training, developing a clear discrimination between cue-on and cue-off states (W= 45, p-value = 0.004) **(Fig. 6C, left)**. Male mice, by contrast, maintained substantial responding during both light on and light off periods throughout training, with no significant trend of divergence between conditions (W=20, p= 0.06) **(Fig. 6C, right)**, despite exhibiting comparable learning trajectories to females as measured by decreasing inter-trial response rates between cue presentation and lever press (sex:day F(6,183) = 0.6540, p-value=0.6868, and increasing success rate (proportion of retrieved reward) over sessions (sex:day F(6,36) = 2.0355, p-value= 0.08606, with a trend of lower success rate in females) **(Figs. S10 and S11)**. We also tested cue discrimination directly as a predictor of the optogenetic effect. We computed a per-animal discrimination index, DI = (rate cue-on – rate cue-off)/(rate cue-on + rate cue-off), from the final three training days, and quantified the effect as log(presses under cue-locked stimulation / mean presses under the no-stimulation and random-stimulation conditions). Across the full cohort, DI predicted the optogenetic effect (Spearman ρ = 0.74, p = 0.006, n = 13; regression β = 1.70, 95% CI [0.49, 2.91], p = 0.010). Because DI covaried with sex, we also examined each sex separately (**Fig. S12**). In females, DI tracked the magnitude of the effect closely (ρ = 0.96, p = 0.003, n = 7; ρ = 0.94, p = 0.017 after excluding the most influential animal), indicating that discrimination carries information about the optogenetic effect beyond sex alone. In males, no association was detectable (ρ = −0.37, p = 0.50, n = 6), although their DI values spanned roughly half the range observed in females (0.01–0.50 vs 0.03–0.90), so a relationship over a wider range cannot be excluded. The DI × sex interaction was not significant (F(1,9) = 2.84, p = 0.126), so we do not claim that the relationship differs formally between sexesMales pressed more during cue-on periods (final 3 days, V = 21, p = 0.031), but maintained substantially higher cue-off responding than females (W = 6, p = 0.035), yielding a lower discrimination index. Because DI was itself lower in males (W = 36, p = 0.035), DI and sex were collinear and explained essentially identical variance (R²= 0.464 for each; AIC 27.83 for each). In the repeated-measures model across all stimulation conditions, the stim × sex interaction remained better supported than stim × DI (F(2,26) = 5.28, p = 0.012, AIC = 510.8 vs F(2,26) = 1.71, p = 0.20, AIC = 520.3).

**Supplementary Table 1.**

| Experiment | Female | Male | Total | Groups |
| --- | --- | --- | --- | --- |
| GI DREADD - Progressive Ratio | 15 | 15 | 30 | Cont GFP (n=16), GI (n=14) |
| GI DREADD - Open Field | 10 | 14 | 24 | Cont GFP (n=7), GI (n=17) |
| GI DREADD - Memory | 8 | 9 | 17 | Cont GFP (n=7), GI (n=10) |
| Food consumption | 6 | 9 | 15 | Cont GFP (n=8), GI (n=7) |
| GQ DREADD - Progressive Ratio | 14 | 14 | 28 | Cont GFP (n=9), GQ (n=19) |
| Stress GQ DREADD - Progressive Ratio | 12 | 9 | 21 | Cont GFP (n=7), Stress GFP (n=6), Stress GQ (n=8) |
| Reward-associated sound conditioning | 18 | 11 | 29 | Cont GFP (n=11), GI (n=18) |
| Fiber Photometry - Reward Discrimination | 4 | 5 | 9 | NA |
| Fiber Photometry - Sensory Response | 2 | 1 | 3 | NA |
| Optogenetic - Progressive Ratio | 7 | 6 | 13 | NA |

**Figure S1.**
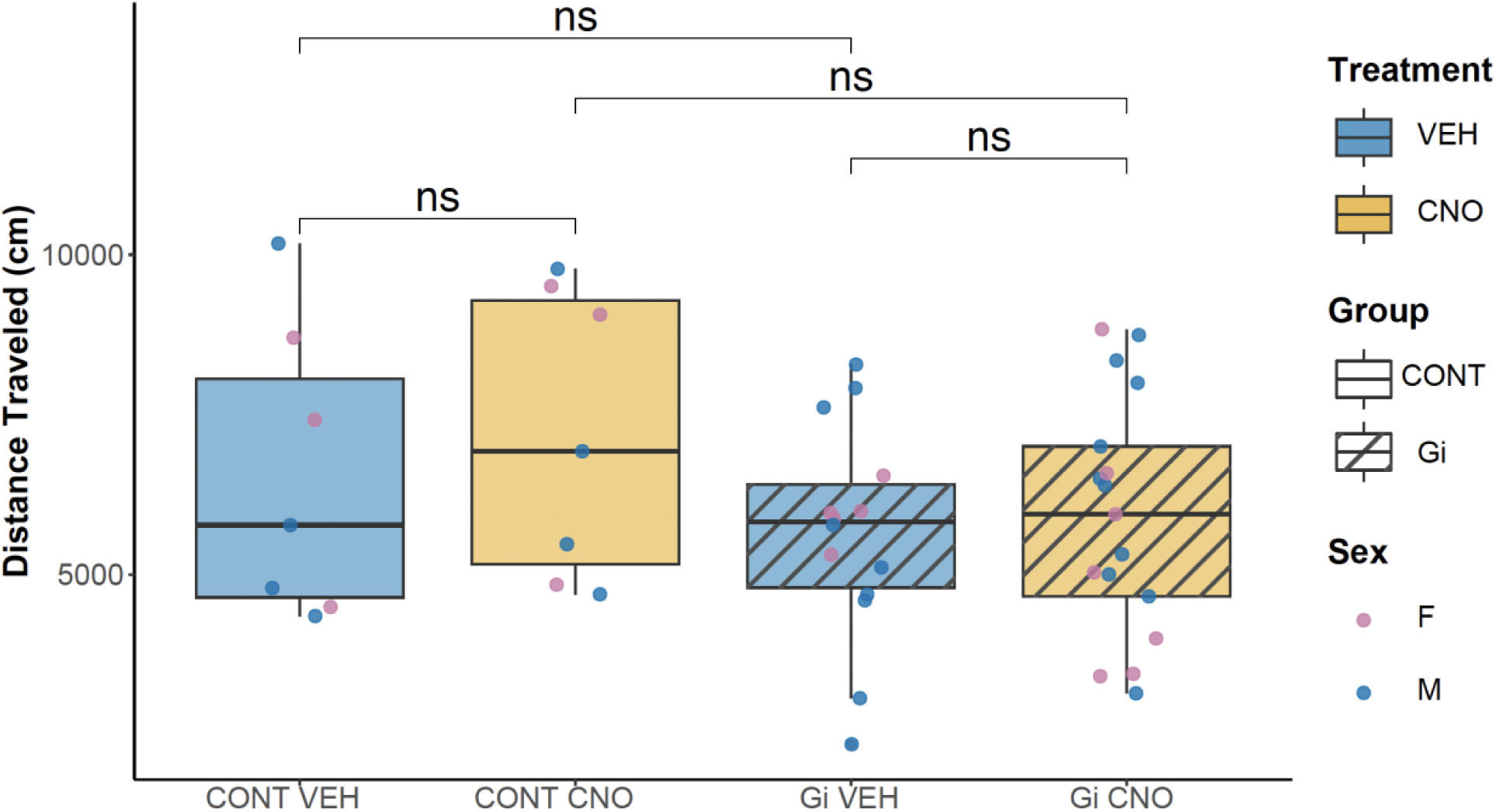
Chemogenetic inhibition of GABAergic ZI neurons does not affect locomotion. Total distance traveled (cm) in the open field for GFP controls and inhibitory DREADD (Gi) mice under vehicle and CNO conditions. No significant differences were observed between any group-treatment combinations, indicating that the reduced pressing after inhibition of GABAergic neurons in the ZI is not attributable to changes in locomotor activity. (Gi: n=18, 10 males, 8 females; GFP: n=17, 9 males, 8 females). ns = not significant.

**Figure S2.**
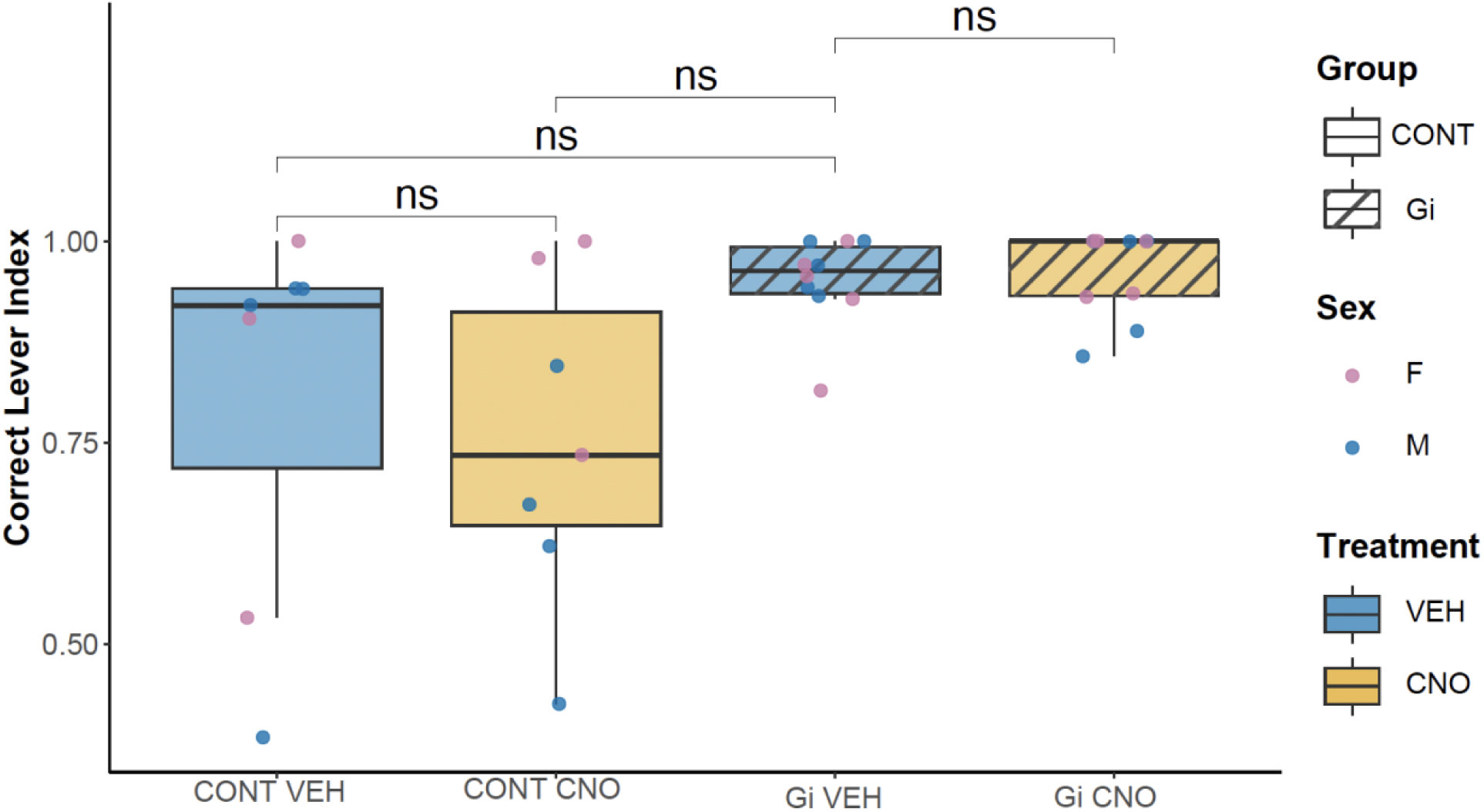
Chemogenetic inhibition of GABAergic ZI neurons does not affect memory. Correct lever index (proportion of presses on the correct (rewarding) lever for GFP controls and inhibitory DREADD (Gi) mice under vehicle and CNO conditions. During training, animals were presented with two levers, only one of which was paired with a food reward; the second non-rewarding lever was present but inactive. No significant differences were observed between any group-treatment combinations, indicating that inhibition of GABAergic ZI neurons does not impair memory. ns = not significant. (GFP: n=7, 4 males, 3 females; Gi: n=10, 5 males, 5 females)

**Figure S3.**
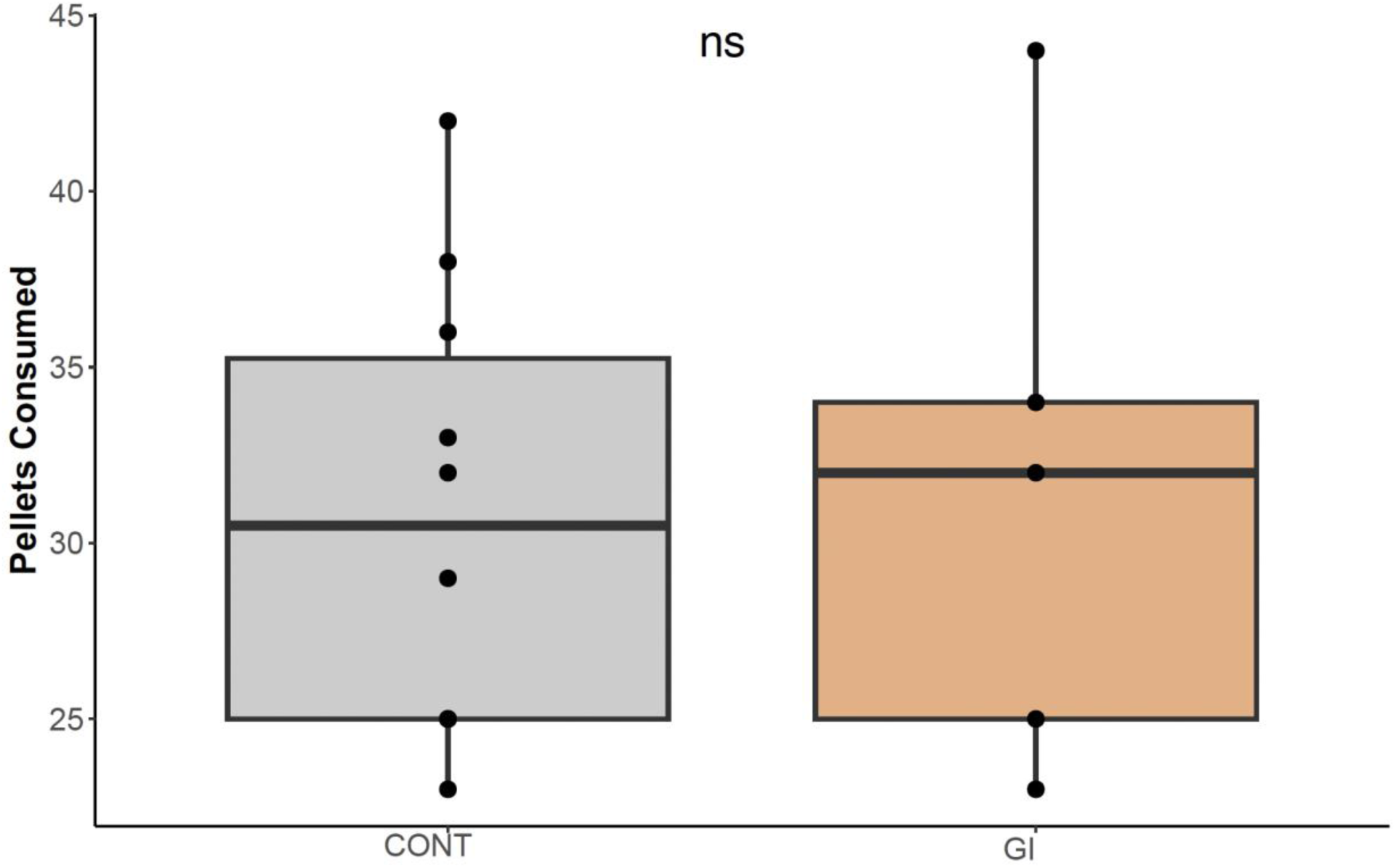
Chemogenetic inhibition of GABAergic ZI neurons does not affect food consumption. Number of pellets consumed by GFP controls and inhibitory DREADD (Gi) mice following CNO administration. No significant difference was observed between groups, indicating that the reduction in breakpoint in PR test is not because the mice become less hungry. (Gi: n=5, 3 males, 2 females, GFP: n=8, 5 males, 3 females).

**Figure S4.**
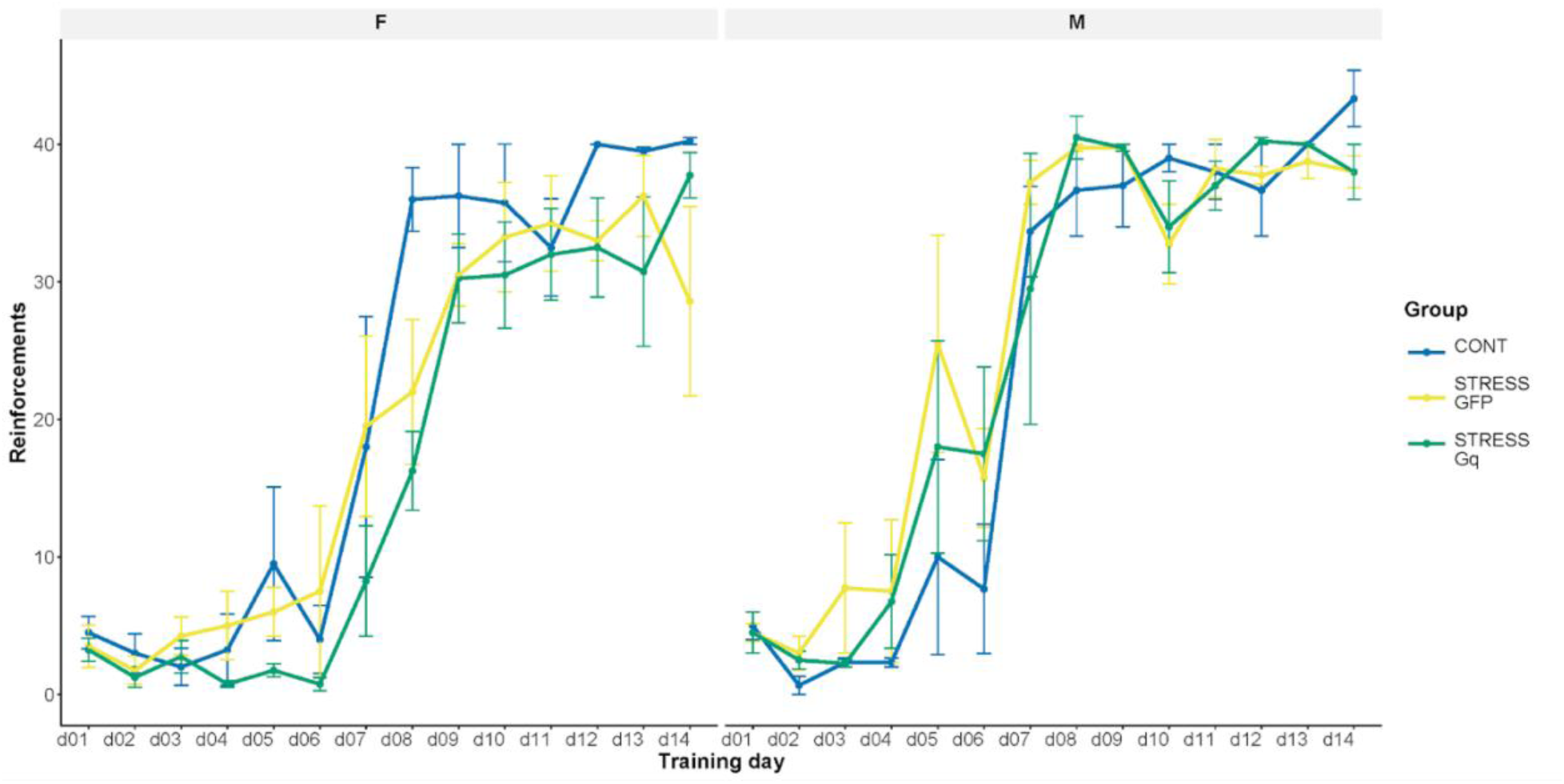
Chronic stress does not impair acquisition during operant conditioning. Number of reinforcements earned across 14 training days for no stress/GFP control, stress/GFP, and stress/hM3DGq groups, representing learning. Separated by sex (females, left; males, right). All three groups acquired the operant response at comparable rates, with no significant differences in reinforcements earned across training days, indicating that stress exposure did not impair task learning.

**Figure S5.**
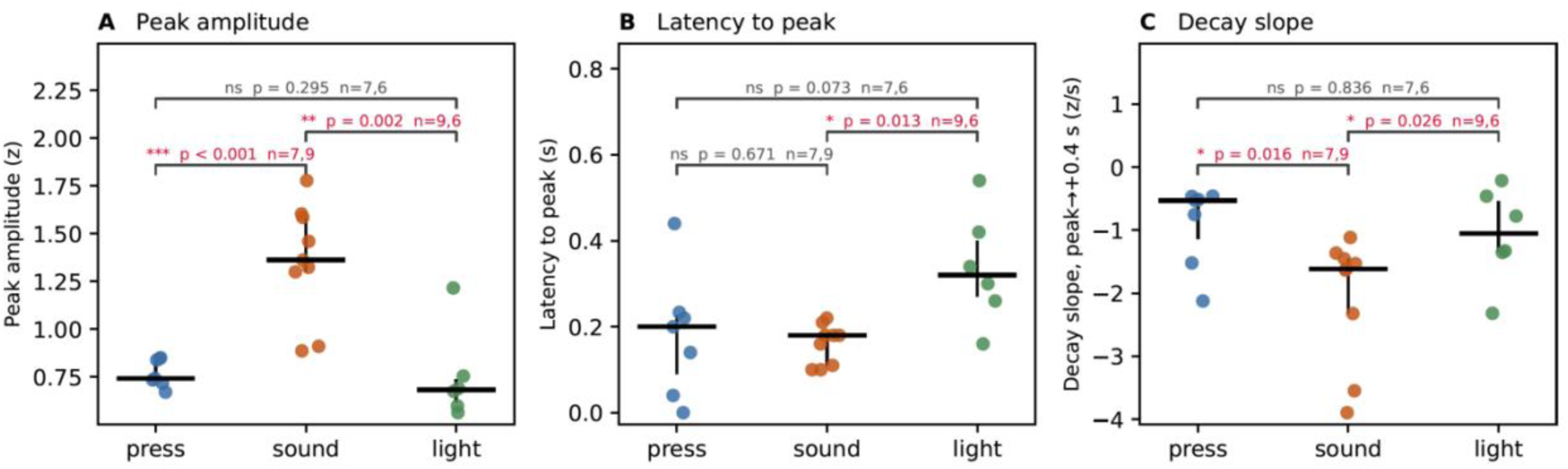
Auditory-cue responses of GABAergic ZI neurons are larger than lever-press and visual-cue responses, with similar amplitude-normalized decay kinetics. Peri-event calcium transients evoked by lever press (blue, n = 7), an auditory cue (orange, n = 9) and a visual cue (green, n = 6) were measured over the same peri-event window (−0.5 to +1.0 s from event onset). **(A)** Peak amplitude of the z-scored ΔF/F transient (Kruskal–Wallis, H = 14.59, p = 0.0007). Auditory-evoked transients were about twice as large as press-evoked (p < 0.001) and light-evoked (p = 0.002) transients. Press- and light-evoked amplitudes did not differ (p = 0.29). **(B)** Latency from event onset to peak response (H = 6.75, p = 0.034). Light-evoked transients peaked later than sound-evoked transients (p = 0.013), and showed a similar, non-significant tendency relative to press (p = 0.073). **(C)** Decay slope, measured over a fixed 0.4 s interval after the peak (z/s; H = 7.71, p = 0.021). Decay was steeper for auditory-evoked transients than for press-evoked (p = 0.016) or light-evoked (p = 0.026) transients. When decay was expressed as the fraction of the peak remaining after 0.4 s, which does not depend on amplitude, event types did not differ (H = 0.45, p = 0.80; not shown), indicating that the steeper absolute decay of auditory responses reflects their larger amplitude rather than faster kinetics. Each dot is one animal, averaged across that animal’s sessions. The horizontal bar marks the median and the vertical line the interquartile range. Brackets show pairwise two-sided Mann–Whitney U tests, with the two group sizes for each comparison.

**Figure S6.**
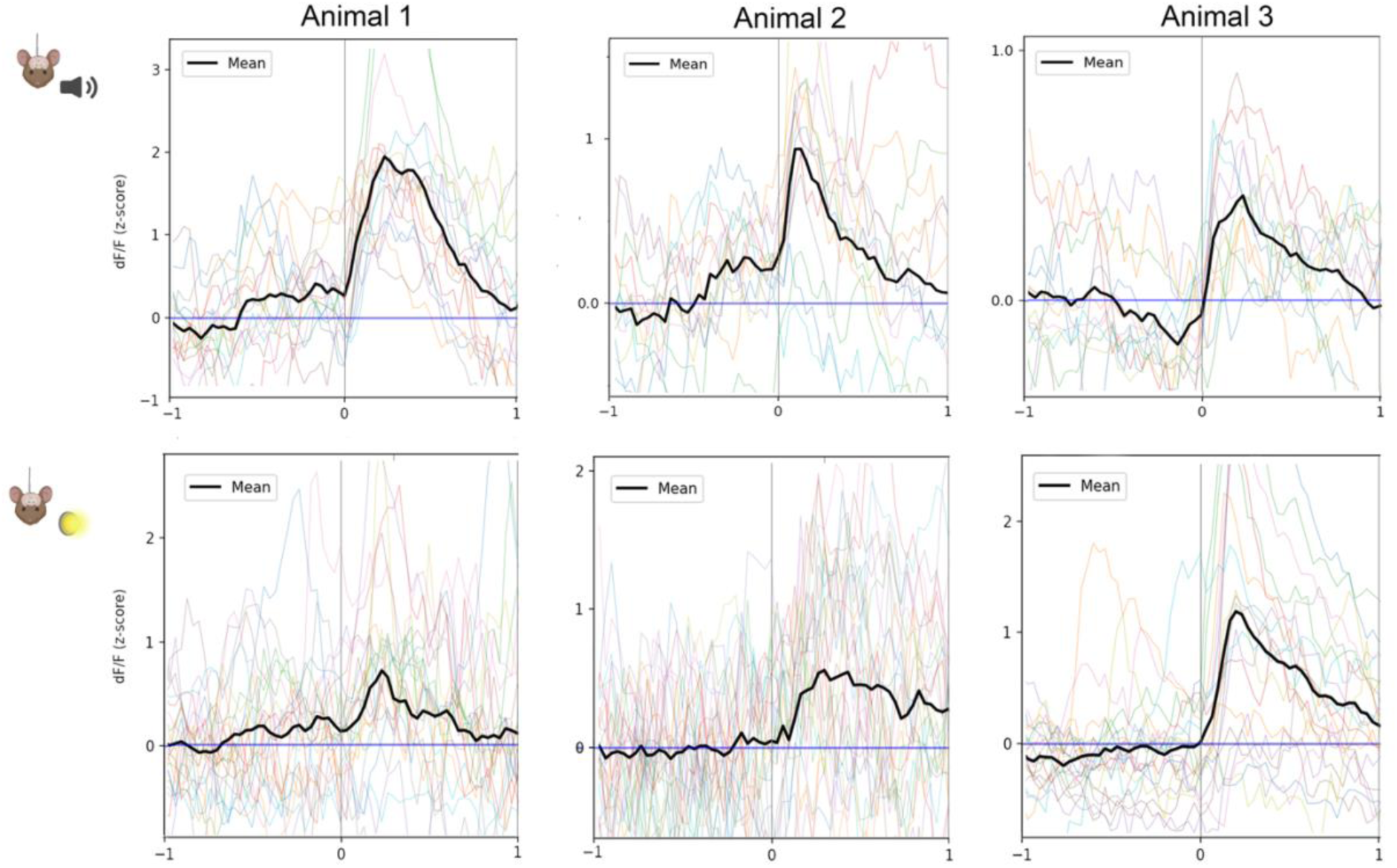
Calcium responses of GABAergic neurons in the ZI of individual animals to auditory and visual stimuli. Z-scored calcium traces (dF/F) from three individual animals in response to auditory (top row) and visual stimuli (bottom row). Individual trial traces (colored lines) and mean response (black line) are shown for each animal, confirming consistent sensory-evoked calcium transients in GABAergic ZI neurons across subjects.

**Figure S7.**
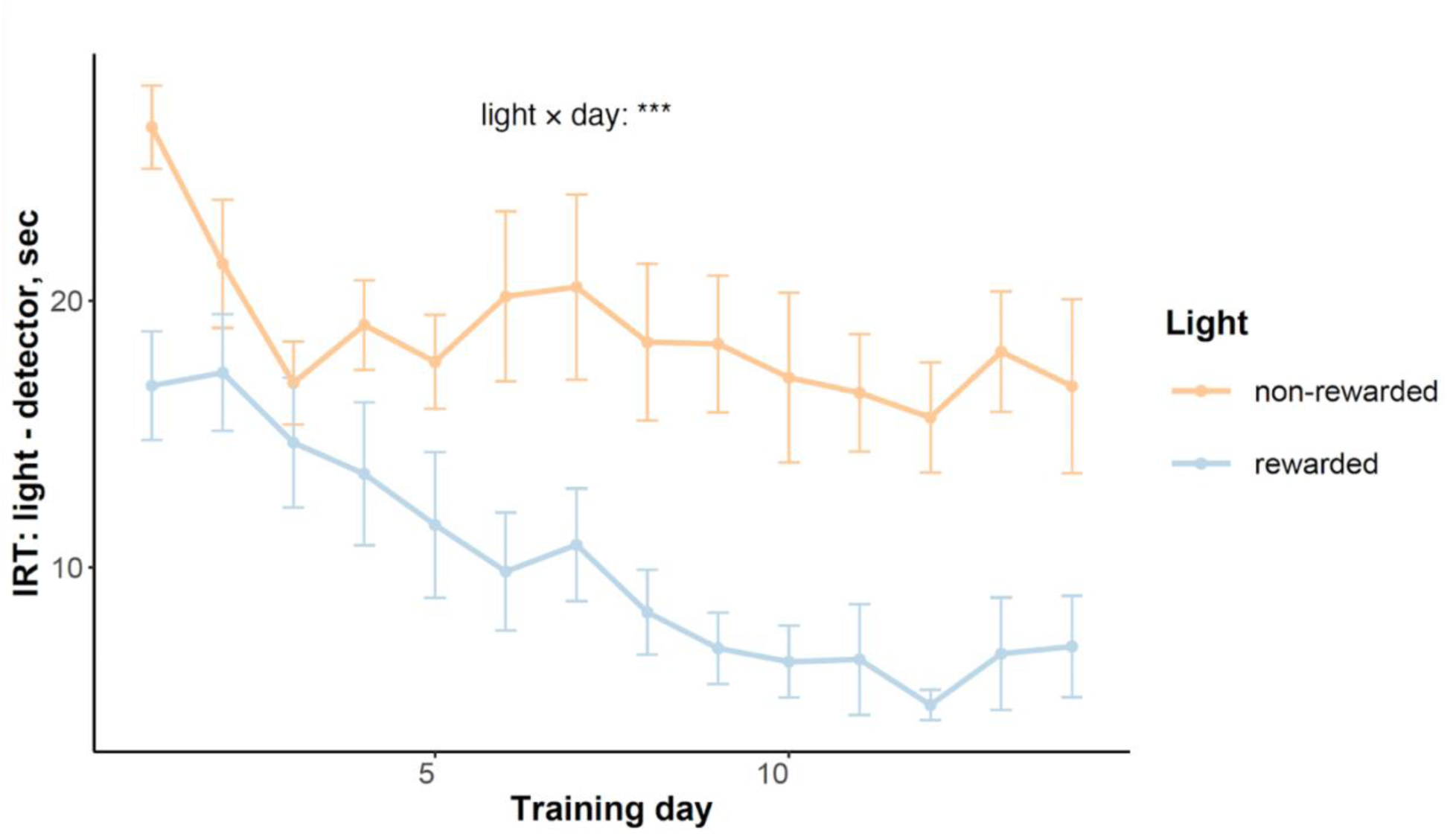
Behavioral evidence of Pavlovian discrimination during training. Inter-response time (IRT) between light onset and detector entry across training days for rewarded (CS+) and non-rewarded (CS-) cues. IRT decreased selectively for the CS+ across training days (significant light x day interaction, p < 0.001), confirming successful acquisition of cue-reward discrimination. (n=9, 5 males, 4 females). Data represented as mean ± SEM.

**Figure S8.**
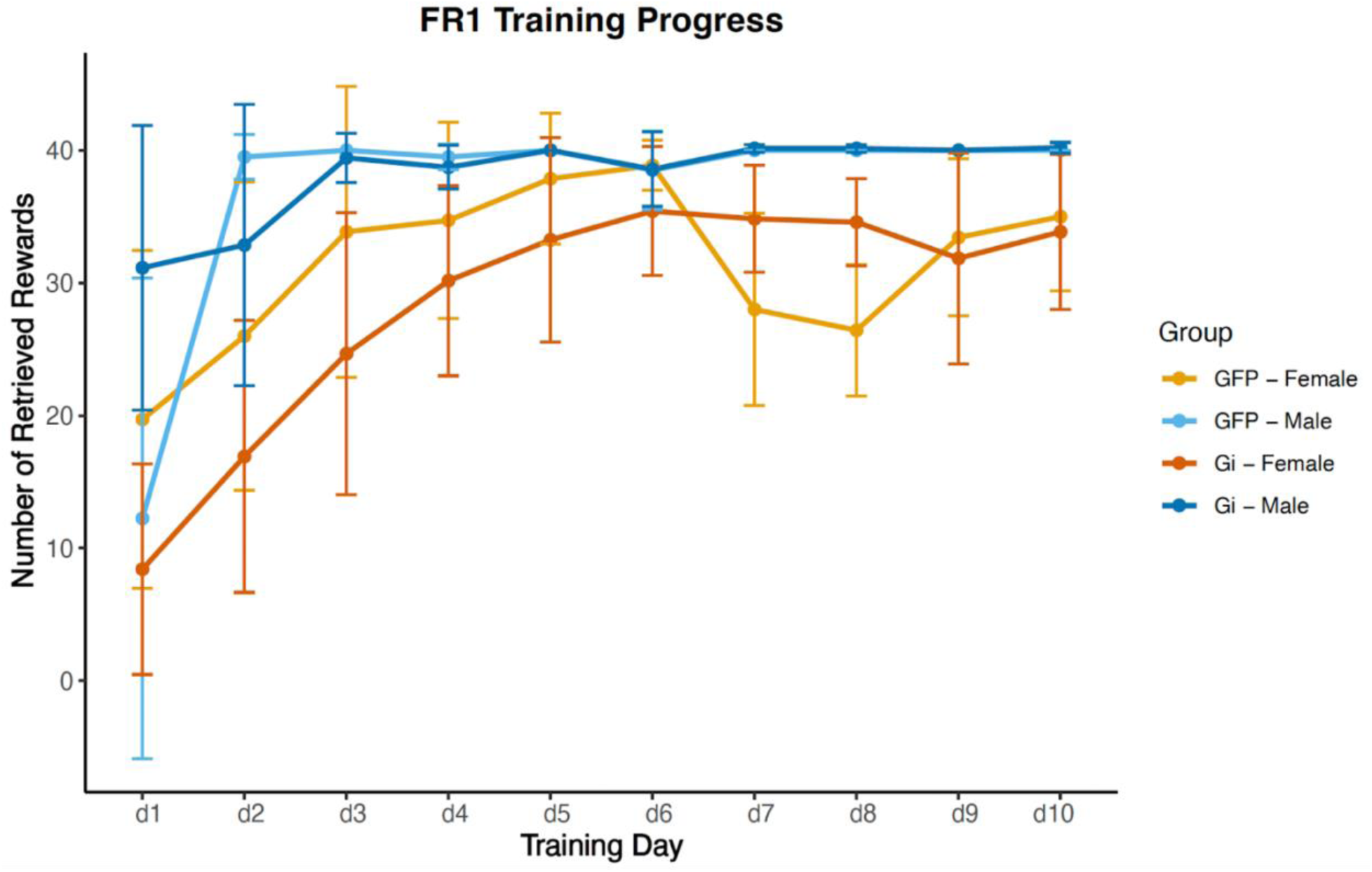
No significant difference in operant training acquisition between Gi DREADD and GFP groups. FR1 training progress showing number of retrieved rewards across 10 training days for GFP and inhibitory DREADD (Gi) groups, separated by sex. (Gi: 7 males, 11 females; GFP: 4 males, 7 females). All groups acquired the operant response at comparable rates. Data represented as mean ± SEM.

**Figure S9.**
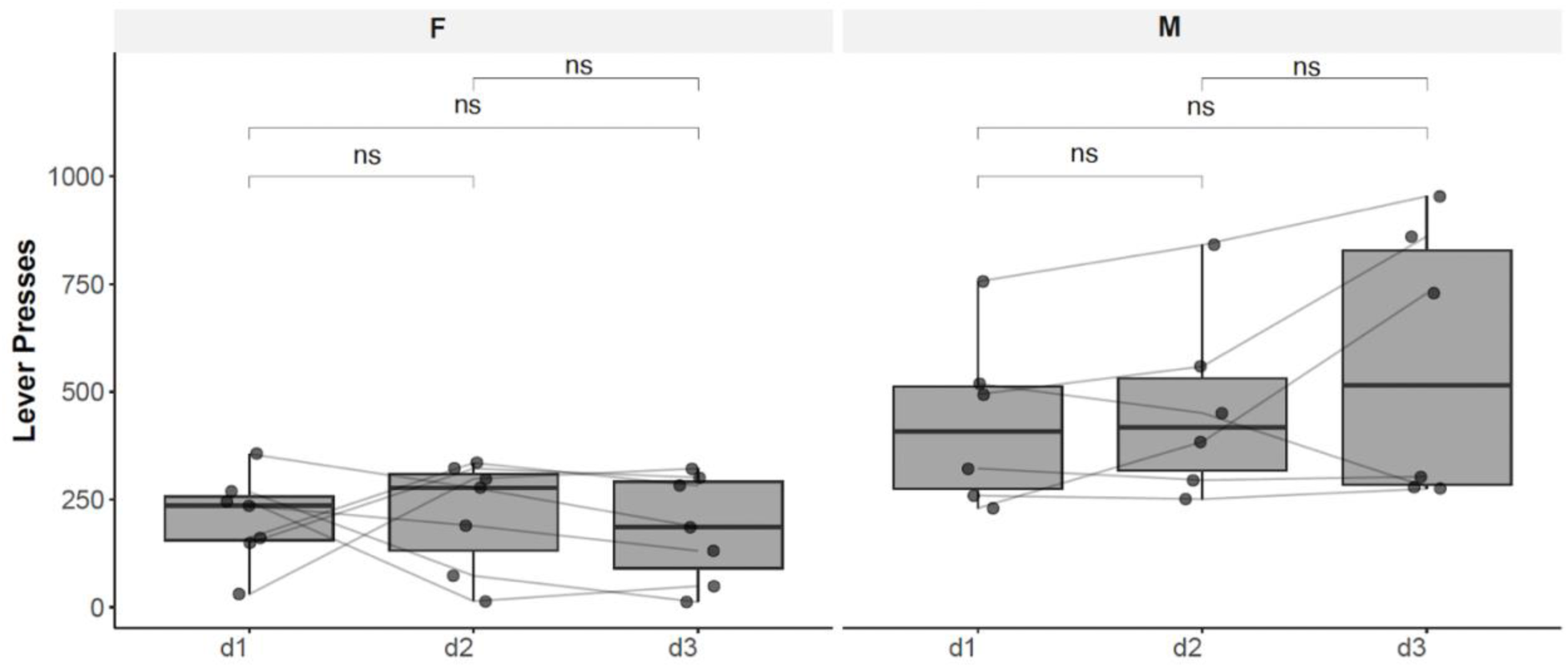
No effect of testing day on PR performance in optogenetics experiment. Total lever presses across three testing days (d1, d2, d3) for females (left) and males (right). No significant differences were observed across days for either sex, ruling out order or practice effects on the optogenetic stimulation results. ns = not significant.

**Figure S10.**
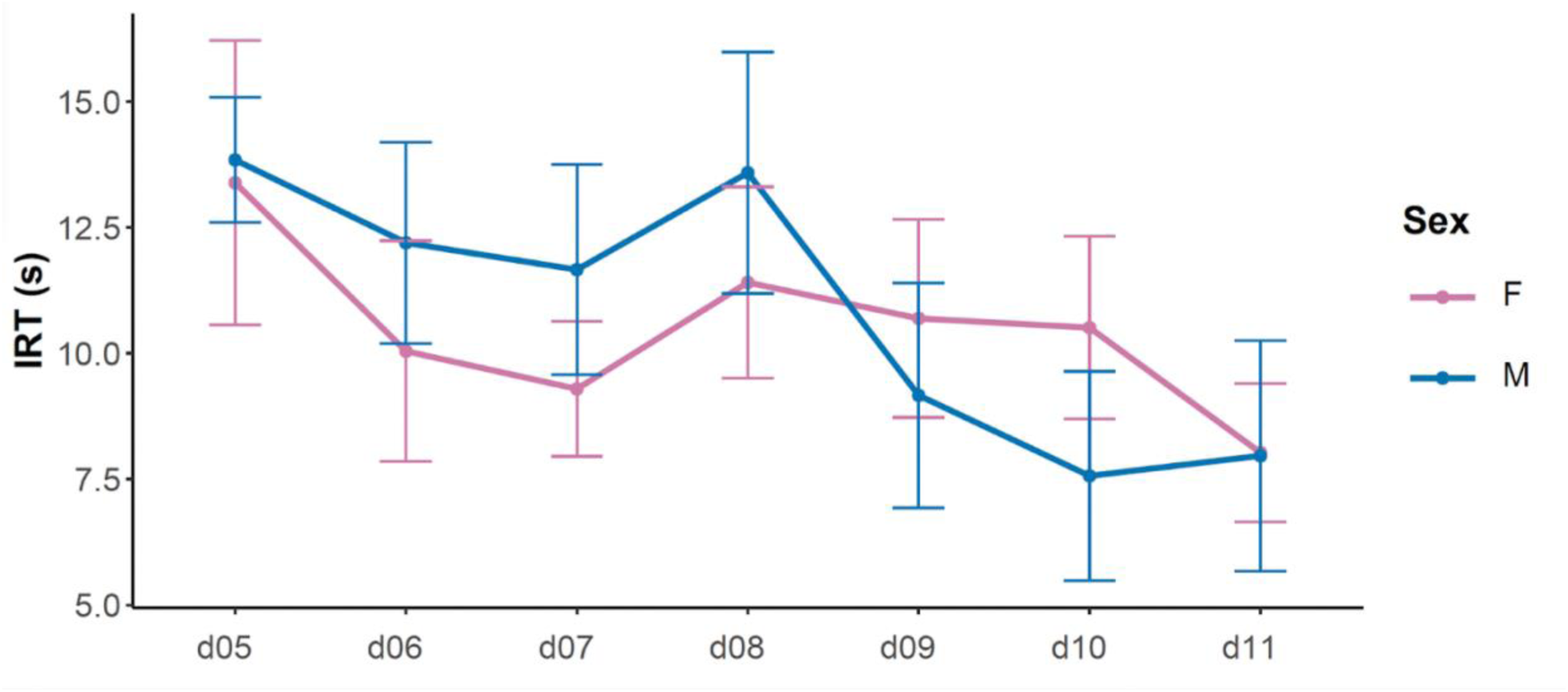
Inter-response time during contingent FR1 training. Inter-response time (IRT, seconds) between light onset and lever press across contingent FR1 training days (d5-d11) for females and males. Both sexes showed comparable decreases in IRT over training, indicating similar learning trajectories.(6 males, 7 females). Data represented as mean ± SEM.

**Figure S11.**
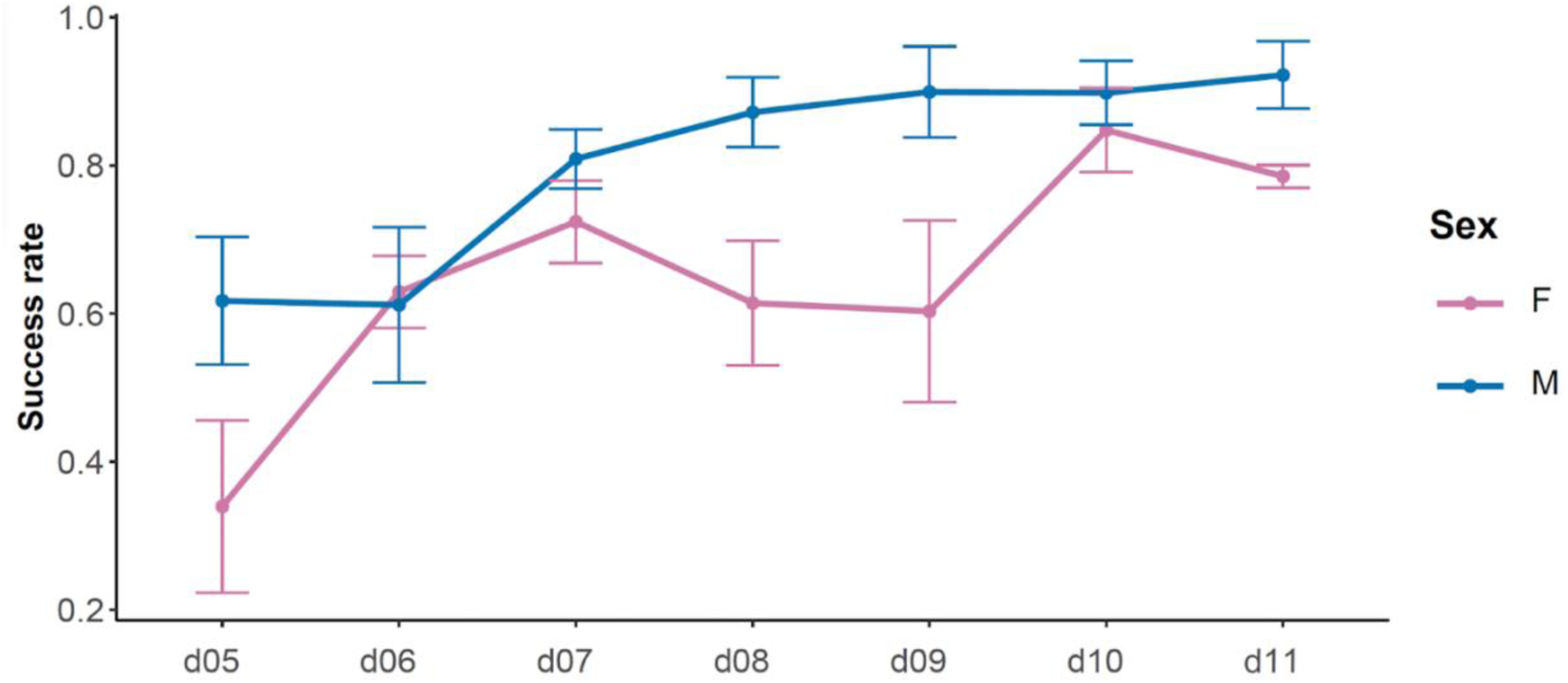
Success rate during contingent FR1 training. Success rate (proportion of retrieved rewards) across contingent FR1 training days (d5-d11) for females and males. Both sexes showed increasing success rates over training, with a trend toward lower success rates in females. (6 males, 7 females). Data represented as mean ± SEM.

**Figure S12.**
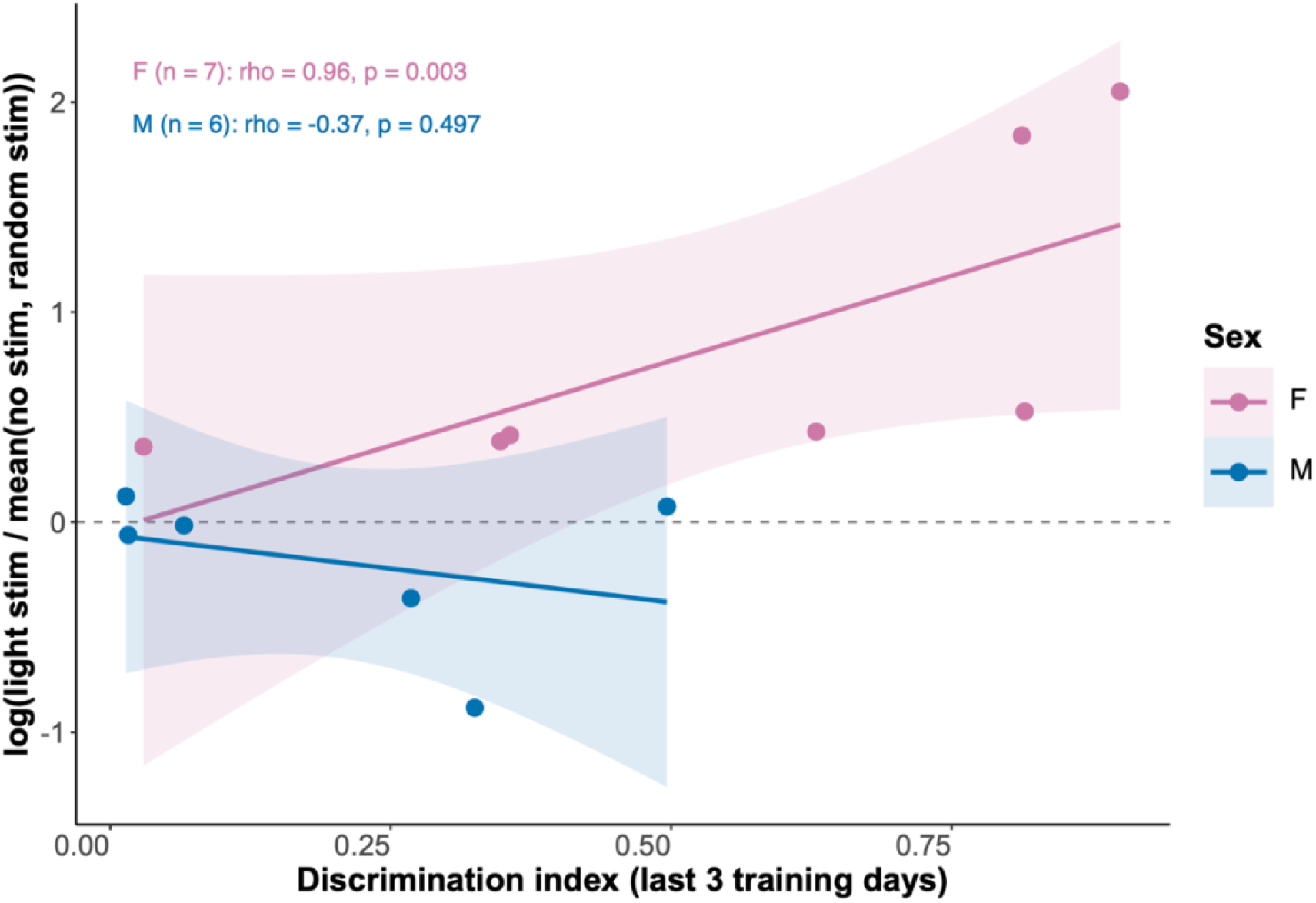
Cue discrimination predicts the optogenetic effect, with the association evident in females. Relationship between each animal’s cue discrimination during training and the effect of optogenetic stimulation of ZI GABAergic neurons on progressive-ratio responding. The discrimination index (DI) was calculated from the last 3 training days as (active − passive) / (active + passive), where active and passive are pressing rates during light-on (lever active) and light-off (lever inactive) periods, respectively. The optogenetic effect is expressed as the log ratio of lever presses during [cue-paired / contingent] light stimulation to the mean of the no-stimulation and random-stimulation sessions. Lines show linear fits with 95% confidence intervals. Within sex, the correlation was significant in females (ρ = 0.96, p = 0.003) but not in males (ρ = −0.37, p = 0.50).

**Fig S13.**
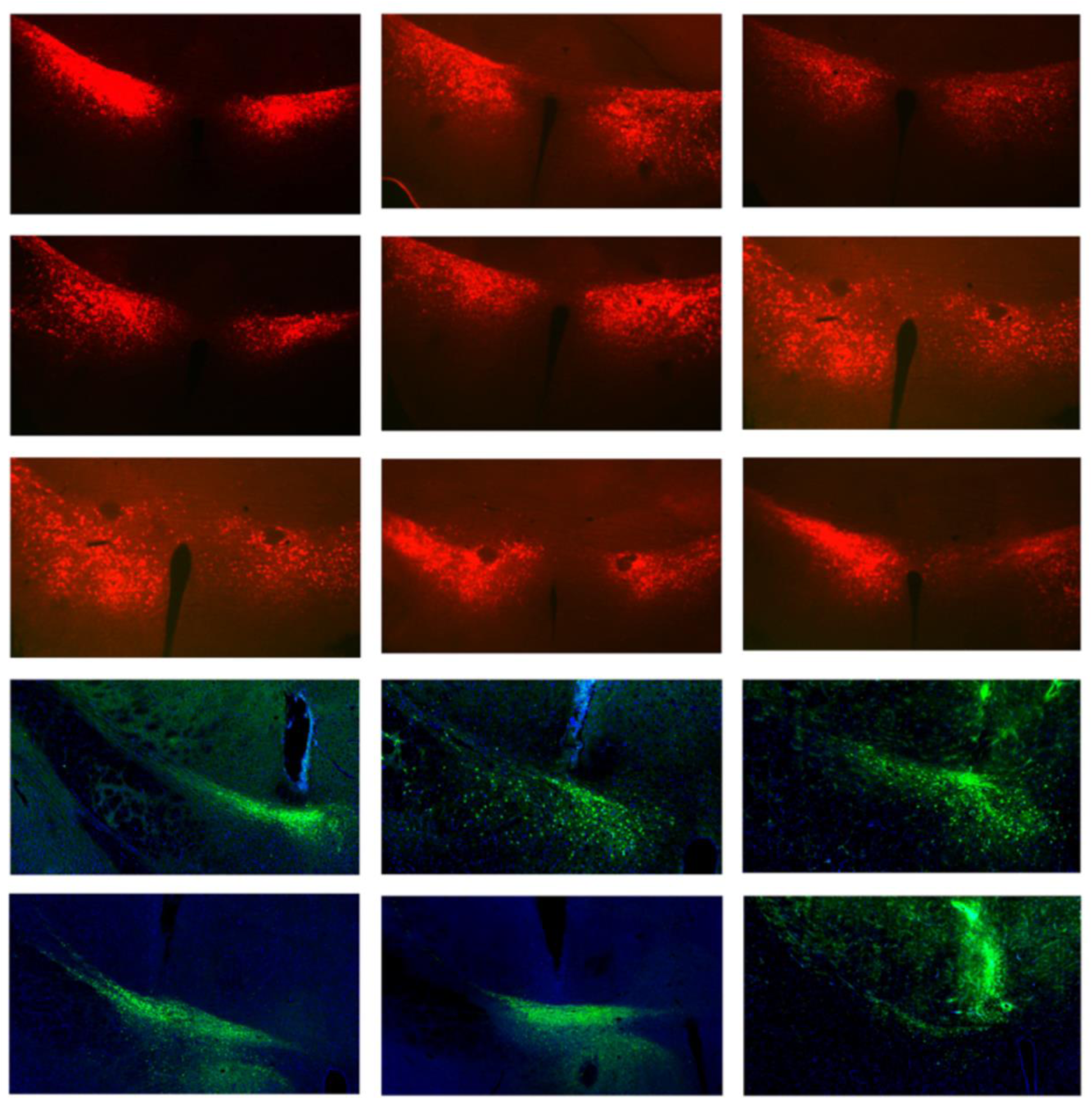
Representative photomicrographs of viral expression in the zona incerta. Rows 1–3: Cre-dependent DREADD expression in ZI GABAergic neurons of vGAT-Cre mice, from nine representative animals. Row 4: GCaMP expression (green) in the ZI, with the optical fiber tract visible above the recording site, used for fiber photometry. Row 5: ChR2 expression (green) in the ZI, used for optogenetic experiments. Sections in rows 4–5 are counterstained with DAPI (blue).

**Fig S14.**
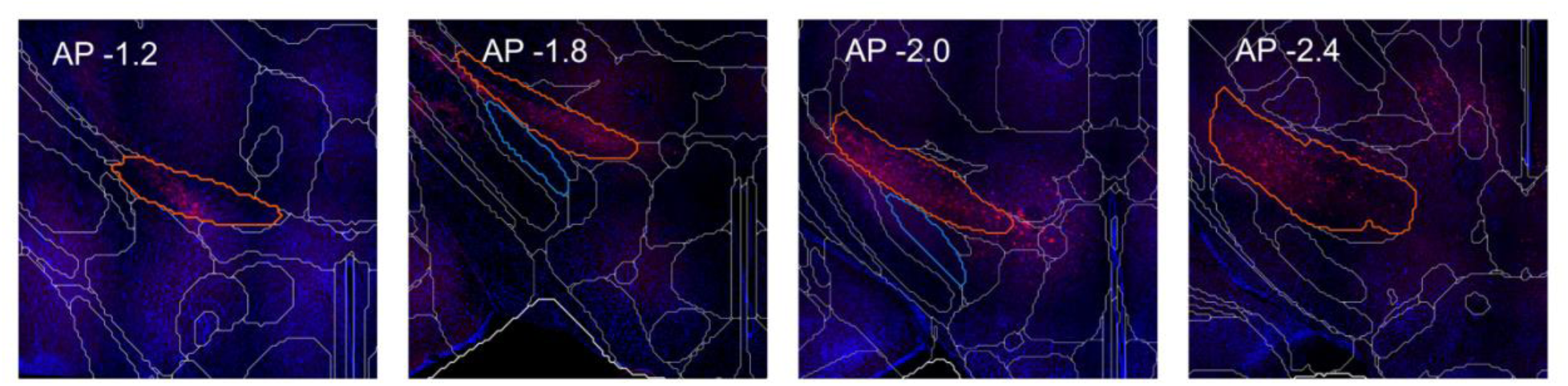
Anterior–posterior extent of viral expression in the zona incerta. Confocal images (20× objective) of coronal sections from one representative animal, showing viral expression along the anterior–posterior (AP) axis of the ZI. Nuclei are counterstained with DAPI (blue). Each section was manually aligned to the corresponding coronal section of the Allen Mouse Brain Atlas; atlas boundaries are overlaid in grey. The ZI is outlined in orange and the adjacent subthalamic nucleus (blue outline) is shown as an anatomical reference and to assess viral spread beyond the ZI. AP values indicate each section’s distance from bregma (mm).

## DISCUSSION

The brain must continuously evaluate which environmental stimuli require attention and action. This computation requires more than reward prediction alone. While dopaminergic signaling is critical for encoding reward value and prediction errors(Coddington & Dudman, 2018; Schultz, 2015), it is insufficient to explain how the nervous system responds to salient cues in stochastic sensory environments and translates those cues into appropriate motivational responses. Here, we first identified GABAergic cells in the ZI as a subcortical cell population that are both necessary and sufficient to modulate effort-based motivation and their stimulation being sufficient to rescue stress-induced deficits in motivation. Next, we determined that these cells function as sensory-motivational integrators. They respond to multi-modal environmental cues, accumulate the significance of these cues over time and are sufficient but not necessary to drive cue-associated motivational output.

Our chemogenetic manipulation of the activity of GABAergic neurons while working for food rewards during progressive ratio testing established that these neurons are both necessary and sufficient for effort-based motivational effort. Inhibition of these neurons reduced progressive ratio breakpoints with large effect sizes, while stimulation enhanced the willingness to press levers for reward. Critically, these effects were specific to motivational state: inhibiting their activity did not alter locomotion, memory, or food consumption, ruling out confounds that the lower breakpoints were a result of movement and memory deficits or satiation. This specificity distinguishes the ZI from nodes such as the ventral tegmental area or nucleus accumbens, where manipulations frequently produce intertwined changes in motor output and hedonic reactivity (Hughes et al., 2020; Kim et al., 2012). The bidirectional control observed suggests that tonic GABAergic ZI activity sets a gain on motivational drive, with the level of ZI GABAergic output scaling the willingness to exert effort for reward (**Fig. 1D, 1E**). This bidirectional control is asymmetric in magnitude. Removing GABAergic ZI output reliably and substantially reduced effort, whereas augmenting it in unstressed, well-trained animals produced a smaller increase that was significant within subjects but did not survive comparison against CNO-treated controls. Motivated responding in food-restricted animals working under a familiar PR schedule may already be near a behavioral ceiling, leaving little dynamic range for a further increase. Consistent with this, activation was markedly more effective where baseline drive had been lowered: in chronically stressed animals, whose breakpoints were significantly depressed, hM3DGq activation produced a large and specific restoration of effort **(Fig. 2E)**. We therefore interpret ZI GABAergic output as setting a gain on motivational drive whose behavioural consequences are most visible when that drive is reduced, either by direct silencing or by chronic stress, and we are careful not to claim symmetric bidirectional control of equivalent effect size.

These findings align with converging evidence that ZI GABAergic neurons promote appetitive and investigatory behaviors across multiple contexts. For example, optogenetic activation of ZI GABAergic neurons strongly promoted hunting of both live and artificial prey, while photoinhibition or deletion of GABA function severely impaired hunting (Ahmadlou et al., 2021; Zhao et al., 2019), indicating that the motivational role of ZI GABAergic signaling extends beyond operant effort to encompass a broad repertoire of goal-directed appetitive behaviors.

With stress-induced deficits in motivation being an endophenotype of depression (Russo & Nestler, 2013; Treadway & Zald, 2011), the rescue of motivational deficits following chronic unpredictable stress via chemogenetic activation of GABAergic ZI neurons carries both mechanistic and translational significance. This result is noteworthy given that a substantial proportion of patients with depression do not respond to conventional monoaminergic treatments (Al-harbi, 2012; Berlim & Turecki, 2007). The rescue of stress-induced deficits in motivation by modulating activity of a GABAergic node suggests the existence of parallel motivational pathways that may be therapeutically accessible when dopaminergic or serotonergic interventions fail. Notably, a GABAergic mechanism for stress-induced anhedonia has precedent. GABAergic, rather than dopaminergic regulation of the VTA is able to mediate depression-like symptoms (Lowes et al., 2021), establishing that non-monoaminergic inhibitory circuits can be causal drivers of motivational impairment under stress. Our finding that ZI activation rescues stress-induced deficits suggests a parallel, non-monoaminergic route for restoring motivation. The ZI’s position as a convergence point for diverse sensory inputs and its long-range inhibitory projections to cortical and subcortical targets (Béreau et al., 2023; Hormigo et al., 2020; Wang et al., 2020; Wilt et al., 2025) make it a particularly attractive candidate for circuit-based interventions in treatment-resistant anhedonia and apathy.

Fiber photometry recordings revealed that GABAergic ZI neurons are not passive sensory relays. These neurons responded to stimuli across auditory and visual modalities, consistent with the known multimodal convergent input architecture of the ZI (Arena et al., 2024; Wang et al., 2020). However, the critical observation was that initially neutral cues acquired the ability to drive differential ZI responses only after Pavlovian conditioning: CS+ cues elicited significantly larger calcium transients than CS-cues after, but not before, training (**Fig. 4D**). This learning-dependent divergence demonstrates that ZI neurons encode the acquired motivational significance of stimulus salience rather than its physical properties. This finding positions the ZI as a node where sensory representations are transformed into motivational signals, a computation that is essential for adaptive behavior. While the amygdala and orbitofrontal cortex are known to encode cue-outcome associations (Herry & Johansen, 2014; Paton et al., 2006; Schoenbaum et al., 2009), the ZI’s role is distinct in that it operates through long-range GABAergic projections capable of directly suppressing activity in downstream targets. Importantly, this role of the ZI in decoding stimulus valence through sensory integration has been corroborated across modalities and behavioral contexts. Visual or whisker sensory deprivation reduced ZI calcium responses to prey and impaired hunting, yet ZI photoactivation largely corrected these sensory deprivation-induced deficits, demonstrating that ZI activity can substitute for degraded sensory input to maintain motivated behavior. In novelty-seeking contexts, optogenetic activation of medial ZI GAD2 neurons dramatically increased interaction duration with novel objects and conspecifics relative to familiar stimuli, elevating arousal levels and deepening investigatory action sequences (Ahmadlou et al., 2021). Strikingly, this discriminative function of the ZI appears to emerge early in development and extends into adulthood. Recent work in pre-weaning pups identified a defined ZI→SST population that integrates multimodal dam-associated cues and selectively responds during mother-infant social interactions (Li et al., 2024). In adulthood, the ZI contributes to sensory discrimination and modulation of sensory information in fear contexts (Venkataraman et al., 2019, 2021). These findings collectively show that the ZI is capable of encoding the learned value of specific stimulus classes and converting them into motivational and state changes.

The optogenetic experiments revealed a surprising and instructive pattern: cue-locked stimulation of GABAergic cells in the ZI enhanced motivation specifically in animals whose behavior was under strong cue control. Female mice, which developed robust discrimination between cue-on and cue-off periods during contingent training, showed significant enhancement of progressive ratio performance when optogenetic stimulation was time-locked to the reward-predictive cue but not when stimulation was delivered randomly. Male mice, who maintained high response rates regardless of cue state, showed no significant effect of any stimulation pattern. This strategic difference is critical for interpreting the optogenetic results. Females developed strong behavioral contingency on the cue, effectively gating their instrumental responding to periods of cue availability. For males, activating GABAergic ZI neurons during cue presentation may not have significantly altered behavior because their pressing was already less contingent on cue information, and they engaged in high levels of operant responding throughout the session regardless of cue presence. Together, these findings demonstrate that activation of GABAergic ZI neurons at the precise moment of a reward-predictive sensory cue is sufficient to boost motivational effort, and that this effect depends not only on the temporal alignment of neural activation with cue processing but also on the strength of the behavioral contingency between cue and action, as revealed by the sex difference in cue-gated responding. This interaction between ZI activation and behavioral strategy suggests that the ZI does not simply add a fixed motivational boost. Rather, it amplifies the signals that are currently controlling action selection. This finding resonates with theoretical frameworks proposing that motivation arises from the interaction of internal states with environmental information (Puglisi-Allegra & Ventura, 2012), and provides a circuit-level mechanism for how this interaction might be implemented. The dependence of ZI effects on behavioral strategy is also consistent with work by Berridge and Warlow (Warlow & Berridge, 2021), who demonstrated that the same circuit node - in their case, the central amygdala - can generate divergent motivational outputs depending on context and the strategy an animal adopts. Our data extend this principle to the ZI, suggesting that the motivational impact of subcortical circuit activation is not fixed but is shaped by the behavioral policy through which an animal engages with environmental cues. The sex difference we observed likely reflects divergent behavioral strategies rather than a categorical sex difference in ZI function per se, as both sexes showed comparable learning trajectories. Nevertheless, this finding highlights the importance of considering individual variation in behavioral strategy when interpreting circuit manipulation experiments and may have implications for understanding sex differences in motivation-related psychopathology.

A seemingly paradoxical finding was that chemogenetic inhibition of GABAergic ZI neurons did not suppress reward-associated conditioned responding. Instead, Gi DREADD mice showed selective increases in pressing during CS+ blocks relative to both acclimation and CS-periods, while GFP controls exhibited a progressive, non-discriminative decline. It is important to be explicit that the PR test and probe test do not measure the same thing. The progressive ratio test measures the effort an animal will expend for reward that is actually delivered, whereas the probe test measures responding that is maintained, and normally extinguished, in the absence of reinforcement. Cue-selective responding of the Gi group reflects the residual behaviour still under the control of the pre-established CS+ association. It is consistent with the involvement of the ZI in the updating and suppression of learned associations in other behavioural domains (Venkataraman et al., 2019, 2021).

An alternative, and not mutually exclusive, account is that cue-driven motivational output is supported by a partly redundant network in which the ZI is one of several nodes. When ZI-mediated drive is attenuated, alternative circuits capable of supporting cue-reward associations, such as amygdala-to-nucleus accumbens, thalamo-striatal, or cortico-striatal pathways (Jennings et al., 2013; Zorrilla & Koob, 2013), may be released from ZI-mediated inhibition or may compensate to maintain cue-directed behavior (**Fig. 5D**). This interpretation also aligns with evidence that VTA GABA neurons can themselves disrupt responding to reward-predictive cues (Wakabayashi et al., 2019), underscoring that cue-driven behavior is supported by multiple inhibitory nodes whose contributions can dissociate under perturbation. This architectural redundancy is consistent with the evolutionary importance of maintaining motivated behavior: a system critical for survival would not depend on a single node. It also suggests that the ZI’s normal role may involve integrating and prioritizing among multiple motivational signals, with its tonic inhibitory output serving to suppress competing or contextually inappropriate responses. ZI inhibition might be delaying the extinction of a conditioned approach response, whereas a redundancy predicts that responding under reinforcement remains cue-selective. When this inhibitory regulation is removed, previously gated cue-reward associations may gain unimpeded access to motor output.

While our work makes a compelling case for the ZI to be included in existing neuroanatomical discussions of effort-based motivation and cue-associated motivational salience, some limitations of our work bear mentioning. Every manipulation and recording in this study used Cre-dependent constructs in vGAT-IRES-CRE mice, so all of the effects we report are attributable to vGAT-expressing GABAergic ZI neurons rather than to the ZI as a whole. Conversely, our data speak only indirectly to the non-GABAergic ZI populations, including the glutamatergic (vGluT2-expressing) neurons and the dopaminergic A13 cells. This caveat is not hypothetical: ZI dopaminergic neurons have been shown to encode motivational vigour during food seeking and to modulate feeding-related behaviour (Ye et al., 2023) so the ZI plausibly influences motivated behavior through more than one cell type, and our design isolates only the GABAergic contribution.

Our chemogenetic and optogenetic manipulations targeted GABAergic ZI neurons as a single population, yet recent work makes clear that this population is molecularly, spatially and functionally fractionated. Emerging evidence suggests molecular heterogeneity within this population that may map onto distinct functional roles (Arena et al., 2024; Wilt et al., 2025). Using single-nucleus transcriptomics together with in situ hybridization, slice electrophysiology and viral circuit mapping,(Kast et al., 2026) resolved four genetically distinct GABAergic ZI populations that differ in spatial position, intrinsic physiology and efferent target. Convergent evidence from transcriptomic profiling of the rostral and medial ZI identified two further non-overlapping subtypes with dissociable functions: Ecel1-expressing rostral ZI neurons projecting to periaqueductal grey that drive self-grooming, and Pde11a-expressing medial ZI neurons projecting to the oral pontine reticular formation that promote sleep-to-wake transitions (Zhu et al., 2025). Functional studies converge on the same conclusion. Activation of medial ZI GAD2 neurons drives novelty investigation and arousal (Ahmadlou et al., 2021), a defined ZI→SST population integrates multimodal dam-associated cues in pre-weaning pups (Li et al., 2024), and ZI GABAergic manipulations have been implicated in hunting (Zhao et al., 2019) and in fear discrimination (Venkataraman et al., 2019). This repertoire of behavior is unlikely to be supported by one homogeneous cell class.

This molecular heterogeneity has direct bearing on the interpretation of our results. Our manipulation of GABAergic cells almost certainly spanned more than one of the transcriptomic subtypes described above and were distributed across the dorsoventral gradient along which those subtypes vary **(Fig. S14)**; our effects are therefore best read as the consequence of shifting aggregate GABAergic tone in the ZI rather than as the signature of any one subtype. The motivational effects we report cannot yet be assigned to a specific efferent pathway, and it remains possible that different components of our findings are carried by different molecular subtypes of GABAergic cells in the ZI. The same inhibition that lowered progressive ratio breakpoints left cue-selective conditioned responding intact **(Fig. 1D**, **Fig. 5C)**, and cue-locked but not temporally random activation enhanced effort **(Fig. 6D)**. Such a pattern could be expected if distinct GABAergic ensembles carry tonic motivational gain and phasic cue-related signals. Resolving this will require intersectional strategies that manipulate and record from the now-defined ZI subtypes, together with projection-specific approaches. We expect such investigations to yield a considerably more nuanced account of ZI contributions to motivation than the population-level one we offer in our study. Identifying the specific downstream targets through which the ZI exerts its motivational effects will be critical for building a complete circuit model of ZI-associated motivational drive.

Our fiber photometry recordings captured aggregate population activity, and single-cell resolution approaches will be needed to determine whether individual ZI neurons encode appetitive salience, aversive salience, or both. The sex difference in the optogenetic experiment warrants further investigation with larger cohorts and additional behavioral protocols to determine whether it reflects a genuine sex difference in ZI circuit function or a difference in cue-utilization strategies that may vary across task contexts. Sex was included as a variable in every experiment and in every statistical model, but the present study was neither designed nor powered to test sex differences, and this constrains how our one sex-divergent result should be read. Statistical models detected no main effect of sex and no sex interaction in the chemogenetic, stress-rescue, photometry or Pavlovian probe experiments, and we accordingly pooled across sexes for visualisation where sex was non-contributory. It bears mentioning that, to our knowledge, the existing literature on ZI contributions to learning, memory and appetitive behavior has not reported sex differences in these functions.

This work establishes the GABAergic zona incerta as a sensory-motivational integration hub that transforms multimodal sensory information into motivational signals. The ZI does not merely relay sensory information but computes motivational salience - the learned significance of stimuli for guiding adaptive behavior. Through bidirectional control of motivational drive, encoding of learned cue valence, state-dependent amplification of cue-driven behavior, and operation within a redundant motivational network, the ZI solves a fundamental problem in adaptive behavior: how the brain translates environmental stimuli into appropriate motivational responses. That activation of this GABAergic node is sufficient to rescue stress-induced motivational deficits opens a new avenue for understanding and potentially treating disorders of motivation through non-dopaminergic circuit mechanisms. Through this work, the “zone of uncertainty” is revealed as a hub where sensory signals acquire meaning and environmental cues are translated into the drive to act.

## MATERIALS AND METHODS

### Animals

vGAT-IRES-CRE mice (Slc32a1tm2(cre)Lowl/J, Strain #:016962, Jackson) were maintained on a C57BL/6J-Aw-J/J B6 agouti background. Mice were 8-10 weeks old at the time of stereotaxic surgery and were group-housed on a 12:12 h light/dark cycle with *ad libitum* access to water. For operant behavioral experiments, mice were food-restricted to maintain 90–93% of their free-feeding body weight. Both male and female mice were included in all experiments, and sex was examined as a biological variable in all analyses. The number of mice used in experiments are listed in **Supplementary Table 1.** All experiments were approved by the Institutional Animal Care and Use Committee of Children’s Hospital Los Angeles in accordance with the National Institutes of Health Guide and followed NIH standards.

### Viral Vectors

The following adeno-associated viral (AAV) vectors were used: AAV5-hSyn-DIO-DREADD-mCherry (inhibitory hM4DGi: Addgene # 44362-AAV5, excitatory hM3DGq: Addgene # 44361-AAV5), AAV5-hSyn-DIO-EGFP (Cre-dependent GFP control: Addgene # 50457-AAV5), AAV5-Syn-FLEX-GCaMP8f (for fiber photometry: Addgene # 162379-AAV5), and AAV5-Ef1a-DIO-hChR2(E123T/T159C)-EYFP (for optogenetics: Addgene # 35509-AAV5). All viruses had titers of ≥1 × 10^12 vg/mL.

### Stereotaxic Surgery

Mice received analgesia (meloxicam (Metacam), 1.5 mg/ml, 1 unit, PO) preoperatively. Next, they were anesthetized with isoflurane (1–2% in O2, induction at 3%), and placed in a motorized stereotaxic frame (Stoelting 51730M). Bilateral injections targeting the zona incerta (ZI) were made at the following coordinates relative to bregma: AP -1.52, ML ±0.74, DV -4.68. A volume of 80 nL of virus was delivered per hemisphere at a rate of 40 nL/min using a Nanoject III. The injection needle was left in place for 10 minutes following injection to minimize backflow and then slowly withdrawn. For fiber photometry and optogenetic experiments, a 200 μm optical fiber (0.37 NA; Neurophotometrics) was implanted unilaterally above the ZI at DV -4.6mm and cemented in place with GC FujiCEM 2. Mice were administered oral meloxicam perioperatively and allowed to recover for 14 days before behavioral testing.

### Chemogenetic Manipulation

Clozapine-N-oxide (CNO; Sigma-Aldrich # C0832-5mg) was dissolved in 2% DMSO sterile saline and administered intraperitoneally at a dose of 1.5 mg/kg 40 minutes prior to testing. Vehicle injections consisted of an equivalent volume of 2% DMSO sterile saline. All mice received both vehicle and CNO injections in a within-subjects with a minimum of 1 day between test sessions.

### Operant Conditioning

Operant conditioning was performed in standard mouse operant chambers (Harvard Apparatus) equipped with two retractable levers, a food pellet dispenser and magazine, a house light, and lights above the levers. Reinforcers were 20 mg grain-based dustless precision food pellets (Bio-Serv). Behavioral events were recorded and controlled by Panlab PACKWIN software.

### Fixed-Ratio (FR) Operant Training

Following food restriction, mice were trained on a fixed-ratio 1 (FR1) schedule in which a single press on the active lever resulted in delivery of one food pellet. Sessions lasted 40 minutes or until a maximum number of reinforcements (40) was earned. Training continued for 14 days. The inactive lever was retracted. Mice were considered to have acquired the operant response when they earned ≥30 reinforcers in a session for at least 3 consecutive days.

### Progressive Ratio (PR) Testing

Following acquisition of operant responding, mice were tested on a progressive ratio schedule in which the response requirement for each successive reinforcer increased according to a geometric progression (defined as n_j_ = 5e^(j/5) − 5) (Killeen et al., 2009). Sessions lasted 4 hours or until the mouse stopped pressing for 5 minutes. The breakpoint was defined as the highest ratio completed (i.e., the last ratio at which the animal earned a reinforcement).

### Chronic Unpredictable Stress Protocol

For the stress rescue experiment, a subset of mice was exposed to a 14-day chronic unpredictable stress (CUS) protocol concurrent with FR operant training (Willner, 2017). The CUS protocol consisted of repeated exposure to varying stressors including water submersion (7 seconds, 3 exposures), forced swimming (3 minutes, 2 exposures), restraint (1h, 3 exposures), cage tilt (3h, 3 exposures), tail suspension (5 min, 2 exposures) and mild footshock (0.05mA, 1 exposure). Stressors were presented in a pseudorandom order such that animals could not predict the next stressor. Non-stressed control animals were handled daily but otherwise remained undisturbed in their home cages. FR1 training sessions began 30 minutes after the end of each stressor. Following CUS and operant training, all groups underwent PR testing under vehicle and CNO conditions as described above.

### Open Field Test

To assess locomotor activity, mice were placed in an open field white plexiglass arena (70 × 70 × 50 cm) and allowed to explore freely for 10 minutes. Total distance traveled (cm) was recorded by an overhead camera and quantified using LimeLight software (ActiMetrics). Mice received either vehicle or CNO 40 minutes prior to testing.

### Memory Test

To evaluate whether chemogenetic inhibition of ZI GABAergic neurons affected memory for the operant contingency, mice were trained on an FR1 schedule in which two levers were available but only one was active (a single press resulted in reinforcement delivery). The correct lever index, defined as the proportion of presses on the previously rewarded lever relative to total lever presses, was calculated following vehicle or CNO administration.

### Food Consumption Test

To assess whether changes in breakpoint reflected alterations in appetitive drive, a food consumption test was conducted. Following CNO administration, food-restricted mice previously trained on the FR1 schedule were placed in the operant chamber. Each visit to the food dispenser resulted in pellet delivery. The total number of pellets consumed was recorded.

### Fiber Photometry

Fiber photometry recordings were performed using Neurophotometrics FP300 system. GCaMP fluorescence was excited at 470 nm (calcium-dependent signal) and 415 nm (isosbestic control). Excitation light was delivered through a 200 um optical fiber patch cord (0.37 NA) connected to the implanted fiber via a ceramic sleeve. Emitted fluorescence was collected through the same fiber. Signals were digitized at 30 Hz and recorded using Bonsai software. The isosbestic signal was used to correct for motion artifacts and photobleaching. Calcium transients were expressed as z-scored ΔF/F and quantified as the area under the curve (AUC) during the 1-second window following CS presentation.

### Delayed-reward FR1 task

To temporally dissociate lever-press and reward-delivery responses, mice were tested on a modified FR1 schedule in which the lever did not retract following a press and reward delivery occurred after a 2-second delay. Fiber photometry recordings were performed throughout the session, and calcium signals were time-locked to lever press and reward delivery events. Timestamps for synchronization were obtained via BNC connectors between the operant chambers and the fiber photometry system. Mice were tested individually.

### Multimodal sensory stimulation

Naïve mice (not previously exposed to operant contingencies) were placed in the operant chamber and exposed to brief auditory (sound of lever retraction) and visual (2-s illumination of the above lever light) stimuli while calcium dynamics were recorded. Stimuli were delivered in a pseudorandom order with a variable inter-trial interval of 20– 40 s.

### Pavlovian Discrimination Conditioning

Mice expressing GCaMP in ZI GABAergic neurons underwent a Pavlovian discrimination protocol in which one visual cue (CS+; light on the right side) was paired with food reward delivery and a second visual cue (CS−; light on the left side) was presented without reward. Each conditioning session consisted of 15 CS+ trials and 15 CS− trials presented in a pseudorandom order. Each CS was presented for 3 seconds, followed by reward delivery (CS+ only) and a variable inter-trial interval. Calcium transients were recorded via fiber photometry before training (first session) and after 7 days of conditioning. Responses were quantified as the area under the curve (AUC) of z-scored ΔF/F during the 1-second window following CS presentation.

### Pavlovian Conditioning with Chemogenetic Inhibition

vGAT-CRE mice expressing inhibitory DIO-DREADDs (hM4DGi) or DIO-GFP control virus in ZI GABAergic neurons underwent Pavlovian conditioning in which an auditory cue (CS+, 6kHz, 70 dB) was paired with reward and a visual cue (CS−) was unreinforced. Following 10 days of conditioning, mice were trained on an FR1 operant schedule for 10 days. Mice then received CNO (0.3 mg/kg, i.p.) 40 minutes prior to a probe test in which CS+ and CS− were presented in alternating blocks without reinforcement. Pressing rate was recorded during each block type (CS+, CS−, and acclimation periods).

### Optogenetic Stimulation Experiment

#### Viral expression and fiber implantation

vGAT-CRE mice received bilateral injections of AAV5-Ef1a-hChR2(H134R)-EYFP into the ZI, and unilateral optical fibers were implanted above the injection sites as described above.

#### Contingent FR1 training

After initial FR1 training (7 days), mice were transitioned to a contingent FR1 schedule in which a visual cue (light) signaled reward availability. The lever was available throughout the session, but lever presses were reinforced only during 10-second light-on periods. Light-off periods served as a signal that reward was unavailable.

#### PR testing with optogenetic stimulation

Following contingent FR1 training, mice were tested on a PR schedule for 1 hour across three counterbalanced sessions: (1) optogenetic stimulation time-locked to the reward-predictive light cue (CS+), (2) temporally random stimulation, and (3) no stimulation. Optical stimulation parameters were: power 5–7 mW; frequency 20 Hz; pulse width 10 ms; duration 10 s. The order of conditions was counterbalanced across mice. Total lever presses served as the primary dependent measure.

### Histology

Following the completion of behavioral experiments, mice were deeply anesthetized with an intraperitoneal injection of a ketamine/dexmedetomidine (Dexdomitor) cocktail (ketamine 100 mg/kg; dexmedetomidine 0.5 mg/kg) and, upon confirmation of loss of pedal and corneal reflexes, transcardially perfused with PBS followed by ice-cold 4% paraformaldehyde (PFA) in PBS. Brains were extracted, post-fixed in 4% PFA for 24 hours at 4°C, and cryoprotected in 30% sucrose in PB for 48 h. Coronal sections of 40 μm were cut on a cryostat and mounted on glass slides. Viral expression (mCherry, eGFP, GCaMP, or ChR2-EYFP reporter fluorescence) and fiber placement were verified by fluorescence microscopy. Animals with off-target viral expression or fiber placements outside the ZI were excluded from analyses.

### Quantification and Statistical Analyses

All statistical analyses were performed using R (version 2022.07.1 Build 554). Data are presented as median and interquartile range unless otherwise noted. A two-way ANOVA with treatment:group and sex as factors and Wilcoxon rank sum test were used to assess effects, with Cohen’s d reported as a measure of effect size.

For the Pavlovian probe test with chemogenetic inhibition, a linear mixed-effects model was fitted with block (CS+, CS−, acclimation), group (Gi vs. GFP), and sex as fixed factors and animal as a random effect, examining pressing rate as the dependent variable. Post-hoc comparisons between block types within groups were performed using Wilcoxon signed-rank tests.

For the optogenetic experiment, within-subjects comparisons of total lever presses across the three stimulation conditions (cue-locked, random, no stimulation) were performed separately by sex using Wilcoxon signed-rank tests. Inter-response times and success rates across training days were analyzed using repeated-measures ANOVAs with sex and day as factors.

Photometry data were processed using custom Python scripts (Python 3.x; NumPy, SciPy, Pandas). The first 1 second and final second of each recording were removed to eliminate edge artifacts, and frames were trimmed to ensure equal samples per channel. The 470 nm and 415 nm signals were deinterleaved. To correct for photobleaching and motion artifacts, a biexponential decay function was fit to the 415 nm isosbestic signal using nonlinear least-squares optimization (scipy.optimize.curve_fit); in cases where the biexponential fit did not converge, a 4th-degree polynomial was used as a fallback. The fitted isosbestic signal was then aligned to the 470 nm signal using iterative robust linear regression. The corrected signal (ΔF/F) was calculated as 100 × (F_470 − F_fitted) / F_fitted, where F_fitted is the robustly fitted isosbestic prediction. ΔF/F values were then z-scored across the entire session (z = (ΔF/F − mean) / SD).

Behavioral events (lever press, reward delivery, or stimulus onset) were identified from TTL digital input transitions (0→1). For each event, a peri-event segment was extracted from the z-scored trace spanning a 0.5–1 s pre-event baseline window and a 1–2 s post-event window (exact windows varied by experiment: 1 s baseline / 2 s post-event for the delayed-reward FR1 task; 0.5 s baseline / 1 s post-event for sensory stimulation and Pavlovian discrimination experiments). Each segment was baseline-normalized by subtracting the mean z-scored ΔF/F of the pre-event window. Individual trial traces were aligned to event onset (time = 0) and averaged within each animal to generate mean peri-event traces. For the Pavlovian discrimination experiment, responses were quantified as the area under the curve (AUC) of z-scored ΔF/F during the 1-second window following CS presentation.

To test whether cue discrimination or sex better predicted the optogenetic effect, a cue-discrimination index (DI) was computed for each mouse from the last 3 days of contingent FR1 training as (presses/min during light-on, lever-active periods − presses/min during light-off, lever-inactive periods) / (the sum of the two). The optogenetic effect was expressed as the log ratio of lever presses under [cue-locked] light stimulation to the mean of the no-stimulation and random-stimulation sessions, normalizing for individual differences in baseline responding. This effect was modelled by linear regression on DI alone, sex alone, and DI and sex together, and the models were compared using AIC and R². The association between DI and the optogenetic effect was additionally assessed with Spearman rank correlations, pooled and within each sex, and a DI × sex interaction term was tested to determine whether the relationship differed between sexes.

For all analyses, significance was set at α = 0.05. Sample sizes for each experiment are reported in the results and figure legends. Both sexes were included in all experiments; where no significant main effect or interaction involving sex was detected, data were pooled across sexes for visualization.

## ACKNOWLEDGEMENTS

We are grateful for animal husbandry and care provided by the veterinarian and staff in the Animal Care Facility at The Saban Research Institute (TSRI). This work was supported by grants to BGD from the National Institutes of Health (Grant Nos. R01MH134873 and R56MH128427), the Department of Pediatrics at the Keck School of Medicine of the University of Southern California, the Developmental Neuroscience and Neurogenetics Program at The Saban Research Institute, and the Child and Brain Development Program of the Canadian Institute for Advanced Research. LK received funding from the TSRI Pre-doctoral Intramural Award. The authors report no biomedical financial interests or potential conflicts of interest.

